# Changes in the functional connectome and behavior after exposure to chronic stress and increasing neurogenesis

**DOI:** 10.1101/2020.06.15.152553

**Authors:** Luka Culig, Patrick E. Steadman, Justin W. Kenney, Sandra Legendre, Frédéric Minier, René Hen, Paul W. Frankland, Catherine Belzung

## Abstract

Addition of new neurons to the dentate gyrus might change the activity of neural circuitry in the areas which the hippocampus projects to. The size of the hippocampus and the number of adult newborn granule cells are decreased by unpredictable chronic mild stress (UCMS). Additionally, one of the notable effects of chronic stress is the induction of ΔFosB, an unusually stable transcription factor which accumulates over time in several brain areas. This accumulation has been observed in many animal models of depression and it could have a protective role against stress, but no studies so far have explored how a specific increase in neurogenesis might regulate the induction and which brain networks might be predominately affected.

We attempted to investigate the role of increasing adult hippocampal neurogenesis on stress-related behavior and the functional brain circuitry involved in mice exposed to UCMS. We used *iBax* mice, in which the pro-apoptotic gene *Bax* can be selectively ablated in neural stem cells, therefore inducibly enhancing survival of newborn neurons after tamoxifen administration. The animals were exposed to UCMS for 9 weeks and treated with tamoxifen in week 3 after the beginning of UCMS. In week 8, they were submitted to a battery of behavioral tests to assess depressive-like and anxiety-like behavior. In week 9, blood was collected to assess basal corticosterone levels, and the animals were sacrificed and their brain collected for ΔFosB immunohistochemistry. Brain-wide maps of ΔFosB expression were constructed and graph theoretical analyses were used to study the changes in brain networks after stress.

UCMS induced negative correlations between the lateral entorhinal cortex and both the hippocampal structures and the nucleus accumbens in the VEH-treated mice, which were not present in other groups. Ranking nodes by degree reveals a strong thalamic-cortical signature in both non-stress (NS) groups. Exposure to UCMS seems to induce activity in thalamic areas and cerebral nuclei, with a different signature in the UCMS TAM group, which seems to completely “disengage” the neocortex and has most of its nodes with the most connections in the thalamic areas.

## 1. Introduction

Major depression is a complex psychiatric disorder with heterogeneous symptoms and a variable course and, as such, is characterized by inconsistent individual responses to treatment. Although there have been advances in identifying both genetic and environmental factors that contribute to the development of depression, the etiology of the disorder is still not completely clear. These factors interact in a complex way, and one of them, prolonged exposure to stress, has been found to be a major risk factor for initiating depressive episodes in some patients (Dumont and Provost, 1999; Hammen, 2005; Paykel, 2003). This concept is the cornerstone of the diathesis-stress model, which explores the way in which predispositions, or diatheses (which can be both genetic and psychosocial), interact with stressors to precipitate a depressive episode in vulnerable individuals (or not in resilient ones) (Willner and Mitchell, 2002).

Implicated in the vulnerability/resilience to stress is ΔFosB, a stable transcription factor which belongs to the Fos family of proteins, but unlike other members of the family, accumulates over time after repeated stress exposure (Perrotti et al., 2008; Vialou et al., 2010). This accumulation has been observed in many animal models of depression, ranging from chronic restraint stress to unpredictable chronic stress (UCMS), the latter being accepted as probably the most valid animal model of depression (Perrotti et al., 2004). Interestingly, one study established a causal link between ΔFosB induction in the nucleus accumbens (NAc), resilience to chronic social defeat stress and response to fluoxetine, a standard antidepressant (Vialou et al., 2010). Namely, it was shown that ΔFosB induction in the NAc is both necessary and sufficient for resilience, and necessary for fluoxetine to elicit its effects on behavior. Most of the research on ΔFosB focused on the specific brain regions such as the nucleus accumbens and the prefrontal cortex (PFC), and while they are key areas implicated in certain symptoms of depression (such as anhedonia), induction of ΔFosB has been observed in many other brain regions implicated in the etiology of depression (Perrotti et al., 2008; Robison et al., 2014; Vialou et al., 2014, 2010). This fact calls for an approach that focuses on multiple brain structures to determine the properties of both long term and large-scale network interactions in the brain.

Such brain-wide mapping of ΔFosB provides an opportunity to overcome some of the limitations of techniques such as post-mortem studies focusing on specific areas (instead of networks) and neuroimaging, which uses only indirect markers of neuronal activation and which suffers from low spatial resolution. These approaches are valuable and have provided many important insights, but a combination of certain aspects of both approaches (cellular resolution of post-mortem studies and network approach of neuroimaging studies) might contribute to a deeper understanding of the complexity of depressive symptoms. This is especially important in light of recent shifts in conceptualization of hypotheses relating to depression, with a trend toward a network-based approach. This network approach proposes that activity-dependent neuronal communication and information processing disturbances may underlie depression, and that antidepressants may function by altering information processing in the affected neural networks (Castrén, 2005; Castrén and Hen, 2013).

The subgranular zone of the hippocampal dentate gyrus (DG) is one of the few areas where the birth of new neurons (i.e. neurogenesis) occurs throughout life (Altman and Das, 1965; Snyder and Cameron, 2012; Spalding et al., 2013), and these adult-born neurons are able to participate in information processing several weeks after their birth (van Praag et al., 2002), making the hippocampus especially interesting in the context of neural plasticity and information processing. Imaging studies have found a reduction of the volume of the hippocampus in depressed patients (Campbell and Macqueen, 2004), and a proposed explanation of this reduction is a stress-induced reduction in neurogenesis (Sheline, 2000), which might be associated with reduced neuronal complexity and connectivity (Castrén, 2005). Animal studies have further explored the link and suggested that that the stress-induced decrease in neurogenesis acts as a contributor to the pathology of depression (Dranovsky and Hen, 2006), a link which is supported by the fact that chronic treatment with antidepressants has an opposite effect, increasing the rate of adult neurogenesis in the dentate gyrus (Czéh et al., 2001; Malberg et al., 2000). This might mean that, in the context of the network-based approach, antidepressants might elicit their effects by restoring compromised information processing through morphological and physiological reorganization of specific neuronal connections in the brain (Castrén, 2005). In light of the information processing hypothesis, these data support the hypothesis that depression is associated with disturbed information processing in crucial neural networks.

As the hippocampus may be involved in mood disorders, particularly through its control of the hypothalamic–pituitary–adrenal (HPA) axis (Mizoguchi et al., 2003; Surget et al., 2011), local changes in the network (addition of new neurons in the DG) might change the properties of the circuit and (via increased plasticity) change the activation of the areas where the hippocampus projects to, directly or indirectly. One of those structures is the previously mentioned NAc, which receives inputs from the ventral hippocampus that regulate susceptibility to chronic social defeat stress (Bagot et al., 2015). In addition, the hippocampus also has widespread cortical and other projections, so it would be interesting to see how the addition of adult-born neurons might change the whole-brain network properties, especially in response to chronic stress exposure.

To explore this question, we utilized heterozygous (HT) *iBax* mice, in which the pro-apoptotic gene *Bax* can be selectively ablated in neural stem cells following tamoxifen injection, therefore inducibly enhancing survival and functional integration of new born neurons in the adult brain (Sahay et al., 2011), as well as wild type (WT) *iBax* mice, in which tamoxifen administration does not affect *Bax* expression. Two weeks before tamoxifen administration, these animals were exposed to a UCMS protocol that in total lasted for 9 weeks, which is a paradigm that elicits a wide array of behavioral and physiological alterations that are consistent with a naturalistic model of depression (Nollet et al., 2012). In week 6, the animals were submitted to the first Cookie test, a behavioral assay with a specific focus on anhedonia, which is related to the NAc. The remaining 4 cookie tests were carried out until week 9. In week 8, the animals were submitted to a battery of behavioral tests to assess depressive and anxiety-like behavior. In week 9, blood was collected to assess basal corticosterone levels, and the animals were sacrificed and their brain collected for ΔFosB immunohistochemistry or used for dissection of brain areas of interest (NAc, medial prefrontal cortex and the amygdala). ΔFosB was used as a marker for chronic neuronal activation, and brain-wide maps of its expression were constructed and graph theoretical analyses were used to study the changes in brain networks after exposure to stress.

## 2. Methods

### 2.1. Animals

Experiments were conducted on two distinct and equally sized populations (HT and WT) of male *iBax* mice, aged 13-22 weeks at behavioral testing (n = 94). HT animals were generated by interbreeding *Nestin CreER^T2^; Bax^f/f^* and *Bax^f/f^* mice as previously described (Sahay et al., 2011), and were maintained on a mixed C57BL/6 and 129/SvEv genetic background. To induce CreER^T2^ mediated recombination of *Bax* in neural stem cells in the adult brain, mice were treated with approximately 54.5 mg tamoxifen/kg body weight, once a day, intraperitoneally for 5 consecutive days. 20 mg/ml tamoxifen (Sigma, T-5648) stock solution was prepared in corn oil and on each day of treatment dissolved to make a 5.5mg/ml fresh solution. For vehicle (VEH), 10ml/kg body weight was injected intraperitoneally, once a day for 5 consecutive days. Unlike HT mice, which carry two floxed *Bax* alleles (*Bax^f/f^*) and one *Nestin-CreER^Γ2^* allele, WT mice do not have floxed *Bax* alleles and thus administration of tamoxifen should not have an effect on the expression of the pro-apoptotic protein BAX and therefore not affect neurogenesis. All animals were group housed and kept under standard laboratory conditions (12/12h light-dark cycle with lights on at 8:30 PM, room temperature 22° ± 2°C, food and water *ad libitum)* in standard cages (42 cm × 27 cm × 16 cm) with shelter for 1 week prior to the start of the experiment. Animal care and treatment was in accordance with the European Union Directive 2010/63/EU.

### 2.2. Experimental design

Animals were divided into eight groups, with 4 of them categorized by environment and treatment: non-stress vehicle (NS-VEH), non-stress tamoxifen (NS-TAM), UCMS vehicle (UCMS-VEH) and UCMS tamoxifen (UCMS-TAM), and each of them further subdivided by genotype: HT or WT (*n* = 11—12 per group). At the start of the experiment, UCMS mice were isolated in individual cages, and non-stressed mice were group-housed with environmental enrichment (shelter and tubes) in a different room, but under the same time and temperature conditions, as noted in 2.1. In the third week of the experiment, animals were treated with either tamoxifen or vehicle and in the eight week went through a series of behavioral tests. This timepoint of tamoxifen administration was selected in order to maximize the observable effect of adult-born neurons during behavioral tests, as there is a 4-6 week window when adult-born neurons exhibit enhanced synaptic plasticity with increased long-term potentiation (LTP) amplitude and decreased LTP induction threshold (Song et al., 2012). In the beginning of the ninth week, CORT was assessed by blood collection 4 days before the sacrifice, with no micro-stressors applied for 24h before the blood sampling. After the blood sampling, the micro-stressors continued to be applied, and were stopped one day before the sacrifice.

Two methods of sacrifice were used in this experiment, and the animals were divided into two groups based on the method. One group of the animals (n = 52) was deeply anesthetized with sodium pentobarbital (100mg/kg, intraperitoneally), transcardially perfused and brains collected for ΔFosB immunohistochemistry (Figure 1A). The other group (n = 42) of animals was sacrificed by CO2 asphyxiation before the brains were removed and dissected, in order to be able to carry out Western blots on regions of interest, which were necessary to do in order to show specificity for ΔFosB, as the antibody used for immunohistochemistry also cross-reacts with an unidentified protein (but not FosB itself), in addition with ΔFosB.

**Figure 1.**
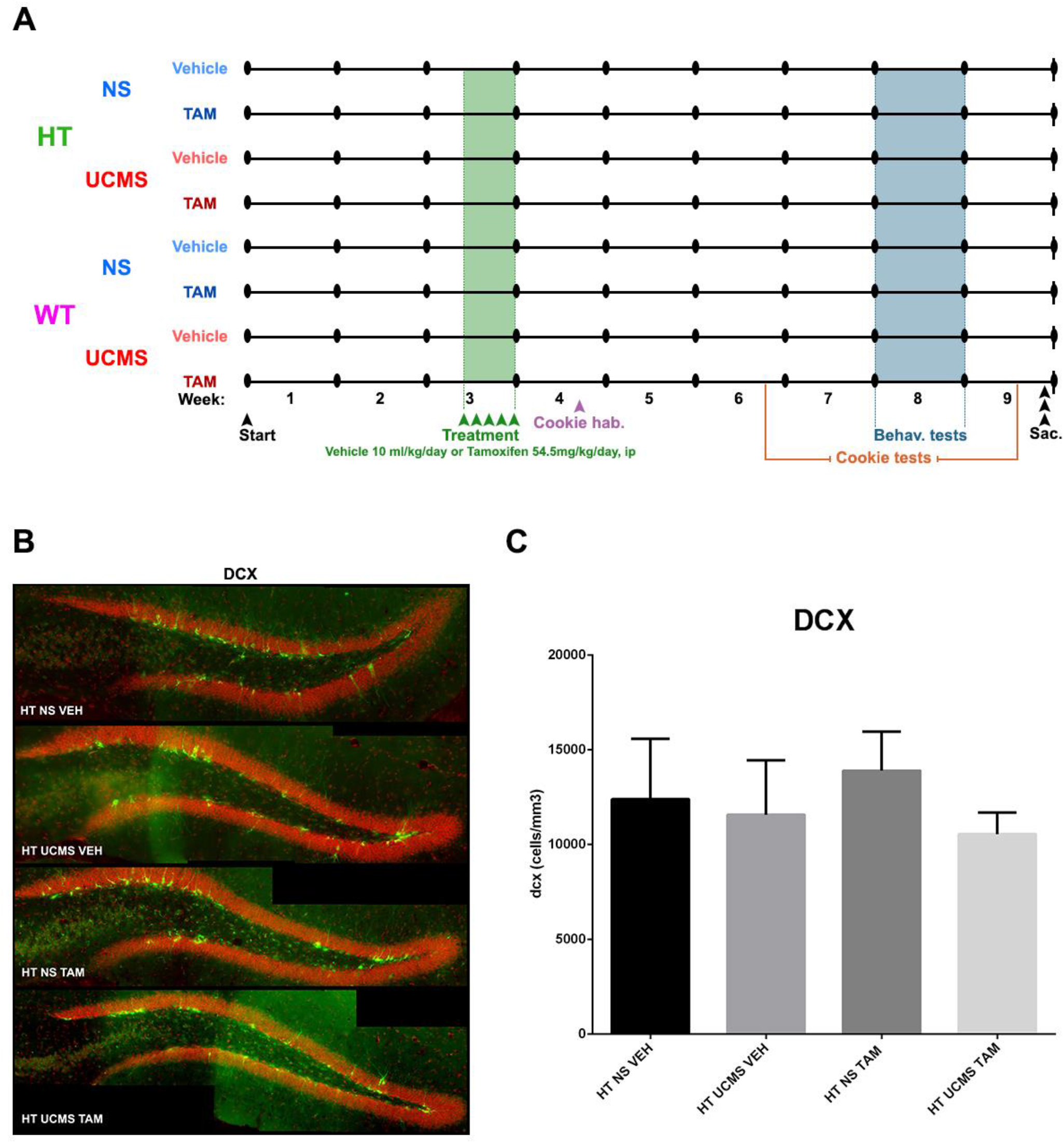
Exposure to unpredictable chronic mild stress (UCMS) and tamoxifen administration had no effect on neurogenesis in the dentate gyrus. **(A)** Schematic representation of experimental design. Mice were either exposed to 9 weeks of UCMS or kept in standard conditions for the same duration of time. Both sets of animals were either treated with 5 daily intraperitoneal injections of TAM or vehicle (corn oil) after two weeks of UCMS. In week 4, animals were familiarized with a cookie for the cookie tests, which were carried out in 5 sessions between week 6 and week 9. In week 8, the mice were submitted to a battery of behavioral tests to assess depressive-and anxiety-like behavior, while in week 9 blood was collected from the animals to assess basal corticosterone levels. **(B)** Representative photomicrographs of immature neuronal marker doublecortin (DCX, green) labeling in the dentate gyrus of the hippocampus. Cell nuclei are counterstained orange with 4’-6-diaminodino-2-phenylindole (DAPI). **(C)** UCMS and TAM had no effect on the number of DCX positive cells, n = 6-7 per group. Data represent mean ± SEM. Abbreviations: HT, heterozygous; WT, wild-type; NS, non-stressed; TAM, tamoxifen; VEH, vehicle; Cookie hab., cookie habituation; Sac., sacrifice

### 2.3. UCMS

The UCMS regimen was used as previously described (Nollet et al., 2013). UCMS mice were isolated in individual cages and subjected to various socio-environmental stressors of mild intensity on a daily basis, according to an unpredictable schedule for 9 weeks. Stressors included removal of sawdust, damping the sawdust, replacing the sawdust with water at 21°C, repeated sawdust changes, tilting the cages by 45°, placing a mouse into a cage that has been occupied by another mouse, predator sounds, restraint stress in small tubes and alterations in the light/dark cycle.

### 2.4. Coat state

The coat state of each animal was assessed at the beginning of each week from Week 1 to Week 9 of the experiment, in order to monitor it as a marker of UCMS-induced depressive-like behavior (Farooq et al., 2012); specifically, as an indirect marker of grooming behavior. The coat state score was calculated as a sum of scores from eight different body parts (head, neck, back, abdomen, forepaws, hindpaws, tail and the anogenital area), each rated on a 0 to 1 scale, with 0 representing good state (smooth and shiny fur), 0.5 representing moderate degradation (fluffy with some spiky patches) and 1 representing bad state (fluffy and dirty fur).

### 2.5. Actimeter

Basal locomotor activity was assessed for 2h in an actimeter during the dark phase of the cycle. All the mice were isolated 36h before the test to avoid a change in activity induced by a novel environment (neophobia). Home cages were placed in a device consisting of a 20 × 20 cm square plane divided into four sections by two photo-beams crossing the plane. The movement of the animal was detected as it crossed the beam, establishing a score representing a realistic measure of the animal’s locomotor activity.

### 2.6. Nest building test

Animals were isolated in individual cages 12h before the start of the test. One square piece of pressed cotton (5 × 5 cm) was placed in each cage at 7:30 AM and the quality of the nest was assessed at three time points (+2.5h, +5h and +24h after placing the nestlet) according to an already described 1-5 rating protocol (Deacon, 2006).

### 2.7. Light-dark box

The device consisting of a light box with transparent sides (20 × 20 × 15 cm), illuminated by a bright light coming from a desk lamp positioned over its center to provide a 1500 lux illumination, and a dark box with the same dimensions and opaque sides. The boxes were connected by a small tunnel (5.5 × 6.5 × 10 cm), which allowed the animals to move freely between them. Animals were placed in the light box, the stopwatch was started and they were observed for five minutes. The latency to enter the tunnel from the light box, total time spent in the dark box and the number of transitions across the tunnel were recorded. An entry to a box was recorded when the animal placed all four paws in the box. The procedure is based on the natural aversion of mice for well-lit areas, and is used as an indicator of anxiety-like behavior (Ducottet et al., 2003).

### 2.8. Splash test

The test was conducted as previously described (Isingrini et al., 2010). Mice were sprayed on their dorsal coat with a 10% sucrose solution in their home cage under red light. The palatability and the viscosity of the solution stimulate grooming behavior. The duration of grooming were recorded during a period of five minutes after the spray.

### 2.9. Tail suspension test

The mice were suspended by the tail using adhesive scotch tape for a duration of 6 minutes, according to a procedure which was previously described (Steru et al., 1985). These six minutes were recorded with a video tracking software (EthoVision, Noldus, Wageningen, The Netherlands), which was also used to analyze these videos. The behavioral measure was the duration of immobility, interpreted as behavioral despair.

### 2.10. Cookie test

This test was carried out in a device containing three aligned compartments with the same dimensions (20 × 20 × 20cm) separated by doors whose opening and closing are managed by the experimenter, as previously described (Surget et al., 2011). Only the colors of the walls and the floor were different between the compartments. Mice were first familiarized with a cookie (Galette St Michel ®) 2.5 weeks before the first cookie test, by placing a small piece of the cookie in the cage of each mouse, in order for the animal to identify its appetizing character. For group-housed NS mice, each animal received one piece of the cookie. On the day of testing, regular food was removed from the cage lid 1 hour before the assay, in order to minimize any potential confounding effects related to differences in satiety. At the time of testing, a small amount of the cookie was placed at the center of the black (third) compartment. The mouse was initially placed in the white (first) compartment of the apparatus. Each session of the test lasted five minutes. When the mouse passed through the first door, it was immediately closed by the experimenter. The cookie consumption (number of bites) was recorded within the 5min test period, as well as the latency to pass the first door, the latency to pass the second door, the latency to smell the cookie, the number of times the animal smelled the cookie and the latency to eat the cookie. Five sessions of testing were performed within 18 days.

### 2.11. Novelty-suppressed feeding test

The test was conducted as previously described (Nollet et al., 2012; Surget et al., 2008). In brief, the test was conducted in a box (33 × 33 × 30 cm) with a sawdust covered floor and under a weak red light. Twelve hours before testing, food was removed from the cages in order to increase the motivation of mice to eat the food pellet (regular chow). At the start of the test, a single food pellet was placed on a white paper (2 × 2 cm) at the center of the box and the animal was placed in a corner of the box. This test induced a motivational conflict between the drive to eat the food pellet and the fear of venturing into the arena. The latency to smell the food, the latency to eat the food and the frequency of smelling were recorded during the testing period of 3 minutes. Immediately after the test, each animal was transferred to its home-cage and the amount of food consumed over 5 min was measured in order to assess differences in appetite, which could act as a confounding factor.

### 2.12. Blood collection and corticosterone level measurement

In week 9, 3 days before the sacrifice, sub-mandibular blood collection was performed for the measurement of plasma corticosterone levels. Plasma was separated from blood samples immediately and stored at −80°C. All blood collection occurred between 10–12h to minimize any potential effects of the diurnal corticosterone variation. The plasma corticosterone levels were then assessed using a commercially available enzyme immunoassay (ELISA) kit (Corticosterone ELISA kit, Enzo Life Sciences) according to the manufacturer’s protocol.

### 2.13 Immunohistochemistry and cell counting

Animals were deeply anesthetized (sodium pentobarbital, 100 mg/kg i.p.) and transcardially perfused with heparinized saline (0.9% sodium chloride, 1000 UI heparin) for two minutes, followed by 80 mL 4% paraformaldehyde in 0.1 M phosphate-buffered saline (PBS, pH 7.4). Brains were collected and postfixed for 2 h in 4% PFA/0.1 M PBS, and then cryoprotected in 20% sucrose/0.1M PBS at 4°C. Coronal sections (40μm) were cut with a cryostat (Leica CM 3050S) and successive sections were placed sequentially in four vials and stored in freezing solution (glycerol, ethylene glycol and PB) at −20 °C for separate immunohistochemical procedures.

For DCX, sections were rinsed in PBS (3 × 10 min) and incubated 48h at room temperature with a goat polyclonal anti-DCX antibody (1:800, sc-8066, Santa Cruz) diluted in PB with 0.1M PB with 0.3% Triton X-100 and 2% normal horse serum (PBH). Sections were rinsed in PB (3 × 10 min) and incubated for 2 h at room temperature with a secondary Alexa Fluor 488 donkey anti-goat antibody (Invitrogen, 1:500). Sections were rinsed in PB again (3 × 10 min) and mounted on gelatin-coated slides and coverslipped with Vectashield HardSet anti-fade mounting medium with DAPI (H-1500; Vector Laboratories).

For ΔFosB labeling, sections were treated in 3% H2O2/50% ethanol for 20 min, rinsed in PBS and incubated with a rabbit monoclonal anti-ΔFosB antibody (1:2500, D3S8R, Cell Signaling) in PBH for 48 hours at room temperature. After three successive rinses in 0.1M PB, sections were incubated for 2 hours with a biotinylated donkey anti-rabbit antibody (1:500, Jackson Immunoresearch) at room temperature. The staining was amplified for 1h with an avidin-biotin complex (Elite ABC kit, Vector Laboratories) and visualized with DAB (Sigma-Aldrich). Stained sections were mounted on gelatin-coated slides, dehydrated in alcohol, cleared with claral and coverslipped.

Cell counting and density measurements for DCX+ cells was performed bilaterally between −0.94 mm to −3.52 mm from bregma (Franklin and Paxinos, 2008), with 5 to 16 sections analyzed per animal. Images were obtained under a ×20 lens with an epifluorescence microscope (Axio Imager.Z2, Zeiss), saved in AxioVision software (Zeiss) and the number of positive cells was counted using Fiji (http://fiji.sc/Fiji). For each section, *z*-stacks of the whole dentate gyrus were acquired and stitched in Fiji with the grid/collection stitching plugin (Preibisch et al., 2009). The granule cell layer area was determined in Fiji and the volume was calculated by multiplying that value with the thickness of the section (40 μm). Cell density was calculated by dividing the number of the cells with the volume (in μm^3^) and multiplying that value with 10^9^ in order for the density to be expressed as number of cells/mm^3^.

For ΔFosB cell counting and density measurements, slide scanned images were processed into individual slices and converted to tiff files. Using Fiji, anatomical labels were manually created for 80 brain regions of interest (ROIs) and for each slice. These ROIs were categorized in major brain subdivisions (hippocampus, thalamus, hypothalamus, cerebral cortex and neocortex), and are listed in Supplementary Table 1. The borders of regions were defined manually according to the Franklin and Paxinos mouse brain atlas (Franklin and Paxinos, 2008). Identification of cells was performed using open-source python tools including scikit-image. Image signal was first inverted and a local contrast enhancement filter was applied which smooths intensity values and adjusts them to either the local max or min depending on which is nearer. A local Otsu filter was applied for object detection followed by a series of binary opening and closing to minimize noise. Objects were then labelled numerically and filtered based on size to remove those with a size too large or small to be a cell. The algorithm was validated using manually counted sections which gave a percent difference of 12% +/− 24% over 54 different regions spread across different animals and slices (Supplementary Table 2). Cell densities were then determined by counting the number of objects detected for a given ROI and divided by the region’s area. For the majority of brain regions three sections were quantified bilaterally and then a mean count was computed for each animal. An example of the procedure is available in Supplementary Figure 2.

### 2.14. Functional network construction and graph theory analysis

Correlation matrices were computed after creating a subset of 66 (out of 80) nodes that were present in all groups of animals. Networks were constructed by thresholding inter-regional correlations in each group of animals by using a cost threshold with an r-value cut-off calculated to keep the top 10% of nodes in each network. The nodes in the networks represent brain regions and the correlations that survived threshold criterion were considered functional connections. Cytoscape software (http://www.cytoscape.org/) was used to visualize the resulting networks, where node size was set proportional to its degree (number of connections) and connection line weights reflect the strength of the correlation. Network measures have been calculated using a custom written R script (https://github.com/jkenney9a/Networks). Definitions and formulae for these graph theory measures have been described in elsewhere (Bullmore and Sporns, 2009; Rubinov and Sporns, 2010; Wheeler et al., 2013). Community detection was conducted with the fastgreedy algorithm, described elsewhere (Yang et al., 2016). For hub identification, nodes which were ranked above the 90^th^ percentile for both degree and betweenness were considered as candidate hub regions for their corresponding network.

### 2.15. Statistics

Given that the assumptions for parametric analyses were not ensured (normality and homoscedasticity), non-parametric statistical tests were performed. Between group effects were assessed by performing the Kruskal-Wallis test and significant main effects were followed by Dunn’s post-hoc test corrected for multiple comparisons. Coat state degradation within each group was assessed by Friedman’s test. Data for Supplementary Figure 1 was analyzed by ANOVA, using ‘environment’ (NS/UCMS) and ‘treatment’ (VEH/TAM) as main factors. When a significant effect of one of these factors or of their interaction was found, the analysis was followed up by a post-hoc Bonferroni’s multiple comparison test. Significance threshold was set at P < 0.05. All data are expressed as mean ± standard error of the mean (SEM).

## 3. Results

### 3.1. Neither exposure to stress nor tamoxifen administration had significant effects on adult neurogenesis in the dentate gyrus

We assessed the levels of neurogenesis in HT mice by immunostaining for the immature neuron marker DCX (Fig. 1B-C). Kruskal-Wallis test did not reveal a significant difference between groups in DCX+ cell density: H_(3,25)_ = 1.79; P = 0.62. The same lack of effect was observed when the hippocampus was segregated into anterior, intermediate and posterior parts and each segment analyzed individually (data not shown). When we segregated the mice into two distinct age-groups, two-way ANOVA revealed an effect of treatment (F_1,10_=11.56, P < 0.01) in the older cohort of mice, and post-hoc Bonferroni’s multiple comparison test detected that TAM treatment induced a significant increase of the density of DCX+ cells in the DG of UCMS animals (UCMS VEH vs UCMS TAM: P < 0.05; Supplementary Figure 1B). Representative pictures of DCX labeling can be seen in Figure 1B.

### 3.2. Effects of stress and tamoxifen treatment on coat state and behavior

In HT animals, coat state deteriorated with time in all groups of animals (Friedman’s ANOVA: P < 0.001) except in the HT NS VEH group (Friedman’s ANOVA: P = 0.58). Kruskal-Wallis test with Dunn’s post-hoc test revealed that exposure to stress deteriorated the coat state in VEH treated animals from week 4 until week 9, with a lack of observed deterioration in week 8 (P < 0.05 for week 5; P < 0.01 for weeks 4, 6 and 7; P < 0.001 for week 9, for HT NS VEH vs HT UCMS VEH), while in TAM treated animals stress exposure deteriorated the coat state from week 2 until week 9 (P < 0.05 for week 2 and 8; P < 0.01 for week 9; P < 0.001 for weeks 3-7, for HT NS TAM vs HT UCMS TAM). No significant effect of treatment has been found between NS groups or UCMS groups (P > 0.05, for HT NS VEH vs HT NS TAM and HT UCMS VEH vs HT UCMS TAM, Figure 2A). In WT animals, coat state deteriorated with time in all groups of animals (Friedman’s ANOVA: P < 0.001) except in the WT NS VEH group (Friedman’s ANOVA: P = 0.11). Kruskal-Wallis test with Dunn’s post-hoc test revealed that exposure to stress deteriorated the coat state in VEH treated animals from week 3 until week 9, with a lack of observed deterioration in week 8 (P < 0.05 for weeks 3-5 and week 7; P < 0.01 for weeks 6 and 9, for WT NS VEH vs WT UCMS VEH), while in TAM treated animals stress exposure deteriorated the coat state from week 2 until week 9 (P < 0.05 for week 3; P < 0.01 for weeks 2 and 6; P < 0.001 for weeks 4-5 and 7-9, for WT NS TAM vs WT UCMS TAM). No significant effect of treatment has been found between NS groups or UCMS groups (P > 0.05 for WT NS VEH vs WT NS TAM and WT UCMS VEH vs WT UCMS TAM, Figure 2B).

**Figure 2.**
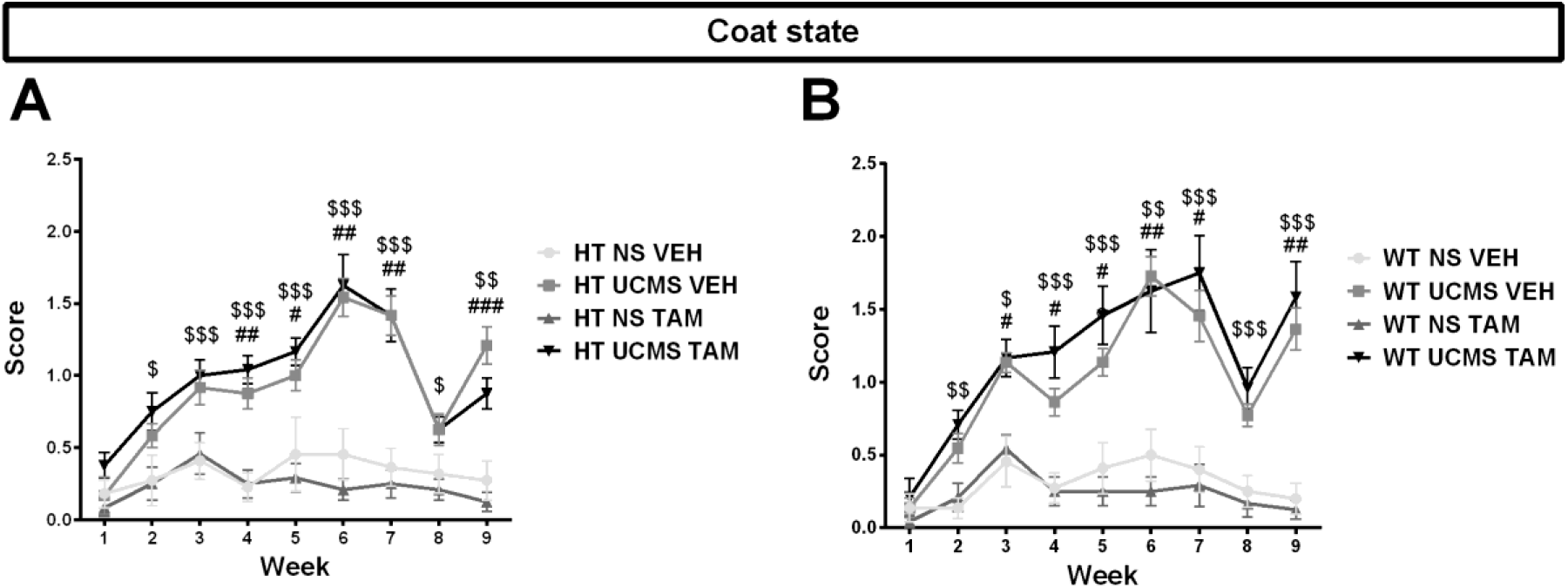
UCMS deteriorated the coat state of both HT and WT animals, while TAM treatment had no effect on coat state. **(A)** In HT animals, coat state deteriorated with time in UCMS mice. Exposure to UCMS induced a deterioration of the coat state in VEH treated animals from week 4 until week 9 (compared to its matched non-stressed group), with a lack of observed deterioration in week 8, while in TAM treated animals stress exposure deteriorated the coat state from week 2 until week 9 (compared to its matched non-stressed group). Treatment with TAM had no significant effect in either the UCMS or NS condition, n = 11-12 mice per group. **(B)** In WT animals, coat state deteriorated with time in UCMS mice. Exposure to UCMS induced a deterioration in the coat state in VEH treated animals from week 3 until week 9, with a lack of observed deterioration in week 8, while in TAM treated animals stress exposure deteriorated the coat state from week 2 until week 9. Treatment with TAM had no significant effect in either the UCMS or NS condition, n = 10-12 mice per group. Data represent mean ±SEM. # P < 0.05 for NS VEH vs UCMS VEH; ## P < 0.01 for NS VEH vs UCMS VEH; ### P < 0.001 for NS VEH vs UCMS VEH; $ P < 0.05 for NS TAM vs UCMS TAM; $$ P < 0.01 for NS TAM vs UCMS TAM; $$$ P < 0.001 for NS TAM vs UCMS TAM

We found no significant effects of UCMS or treatment on locomotor activity in both HT (*H*_(3,46)_ = 1.03; *P* > 0.05, Figure 3A) and WT animals (*H*_(3,42)_ = 0.51; *P* > 0.05, Figure 3B), minimizing the possible confounding effects of a difference in baseline locomotor activity on other behavioral tests.

**Figure 3.**
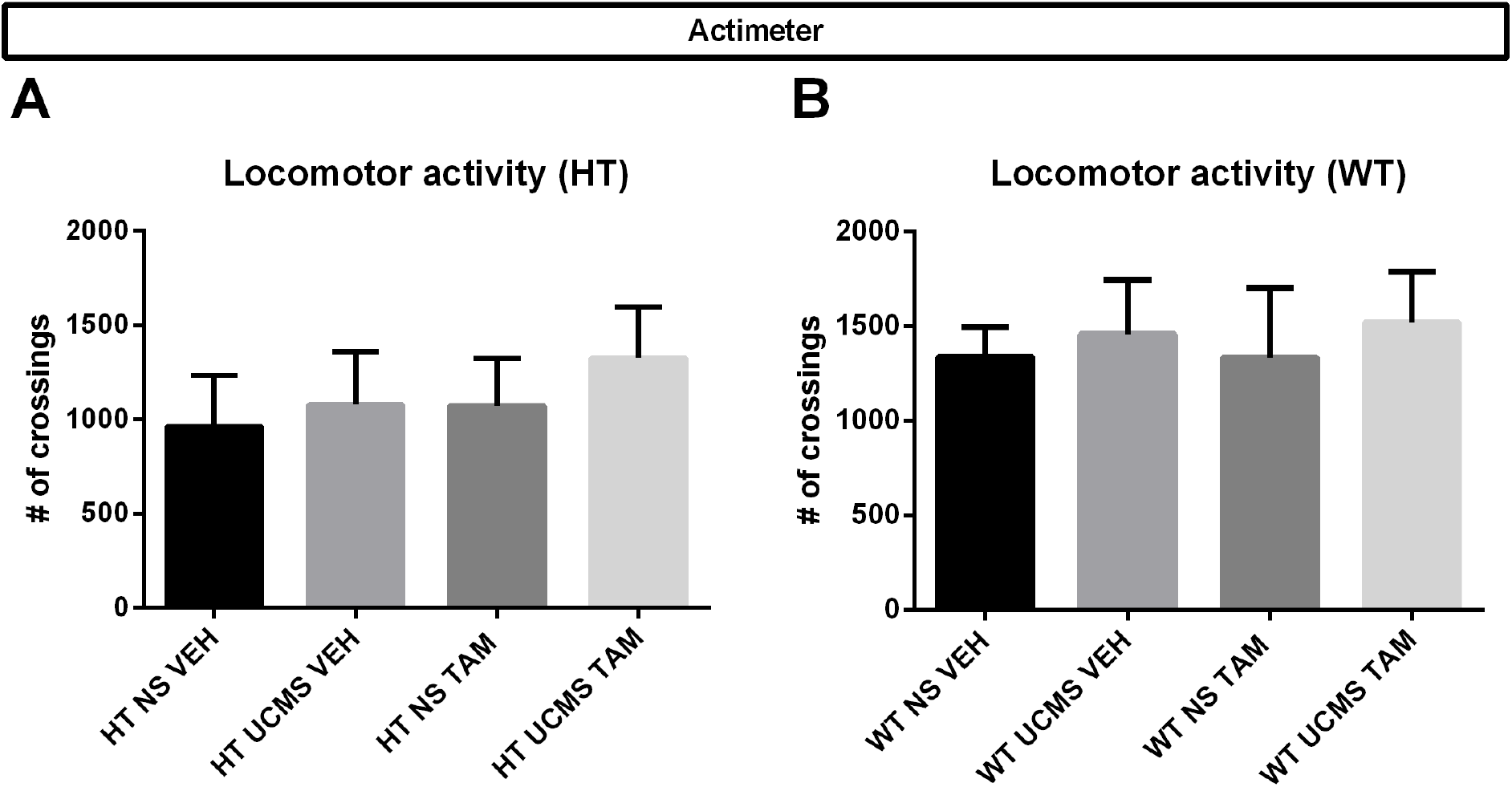
Neither UCMS nor TAM had a significant effect on locomotor activity of HT and WT animals. We did not observe any differences in locomotor activity between **(A)** HT groups and **(B)** WT groups, n = 10-12 mice per group. Data represent mean ±SEM.

Kruskal-Wallis test revealed a difference between HT groups in the nest building test 2.5 h after placement of the nest building material *(H*_(3,46)_ = 8.44; *P* < 0.05) and 5 h after the placement (*H*_(3,46)_ = 10.24; *P* < 0.05), but not 24 h after. No differences were found between the WT groups. Exposure to UCMS improved the nest building score in TAM-treated mice (*P* < 0.05, Figure 4C) at the +5 h time point, an effect which was not observed in VEH-treated mice.

**Figure 4.**
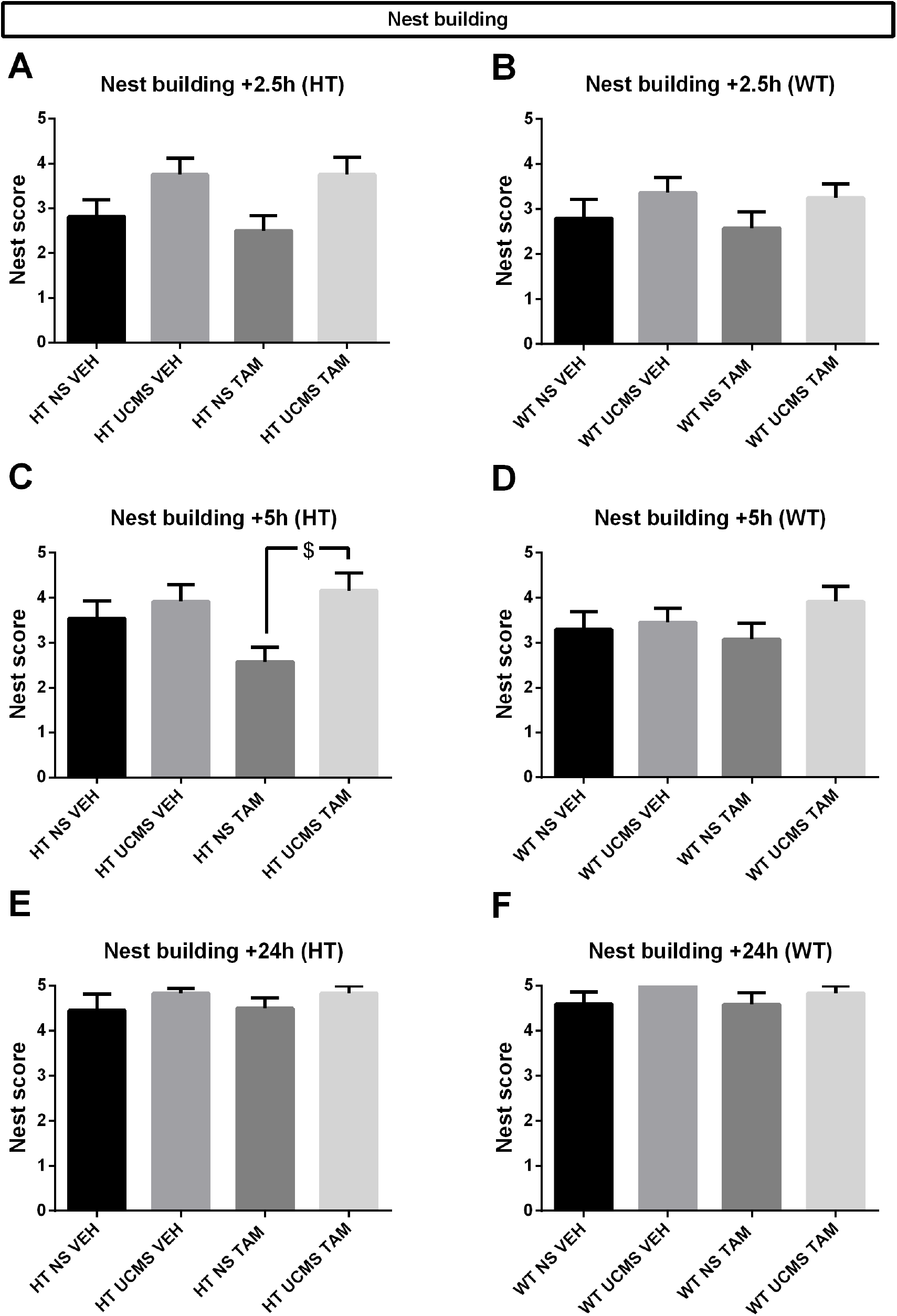
Exposure to UCMS in HT TAM-treated mice improved nest building scores at the +5h time point, while no significant effects were observed in WT animals. **(A)** Exposure to UCMS improved the nest building score in TAM-treated HT mice at the +5 h time point, an effect which was not observed in VEH-treated mice, n = 11-12 mice per group. **(B)** We did not observe any significant differences in nest building scores between WT groups, n = 10-12 mice per group. Data represent mean ±SEM. $ P < 0.05 for NS TAM vs UCMS TAM

In the light-dark box, we found differences between HT groups in time spent in the dark box (*H*_(3,46)_ = 12.52; *P* < 0.01), frequency of entering the light box (*H*_(3,46)_ = 13.96; *P* < 0.01), time in the light box (*H*_(3,46)_ = 14.99; *P* < 0.01) and in the frequency of entering the dark box (*H*_(3,46)_ = 12.33; *P* < 0.01). In HT animals treated with VEH, exposure to UCMS induced a tendency of decreasing time spent in the dark box (p = 0.085, Figure 5A). Similarly, exposure to UCMS in VEH-treated animals increased the frequency of entering both the light box (*P* < 0.05, Figure 5C) and the dark box (*P* < 0.05, Figure 5G). Exposure to UCMS increased the time spent in the light box for TAM-treated animals (*P* = 0.05, Figure 5E). Kruskal-Wallis test revealed a difference between WT groups in the time spent in the dark box (*H*_(3,44)_ = 8.01; *P* < 0.05) and in the time spent in the light box (*H*_(3,44_) = 10.07; *P* < 0.05), but Dunn’s post-hoc tests with corrections for multiple comparisons failed to observe any (a priori chosen) significant pairwise differences between groups.

**Figure 5.**
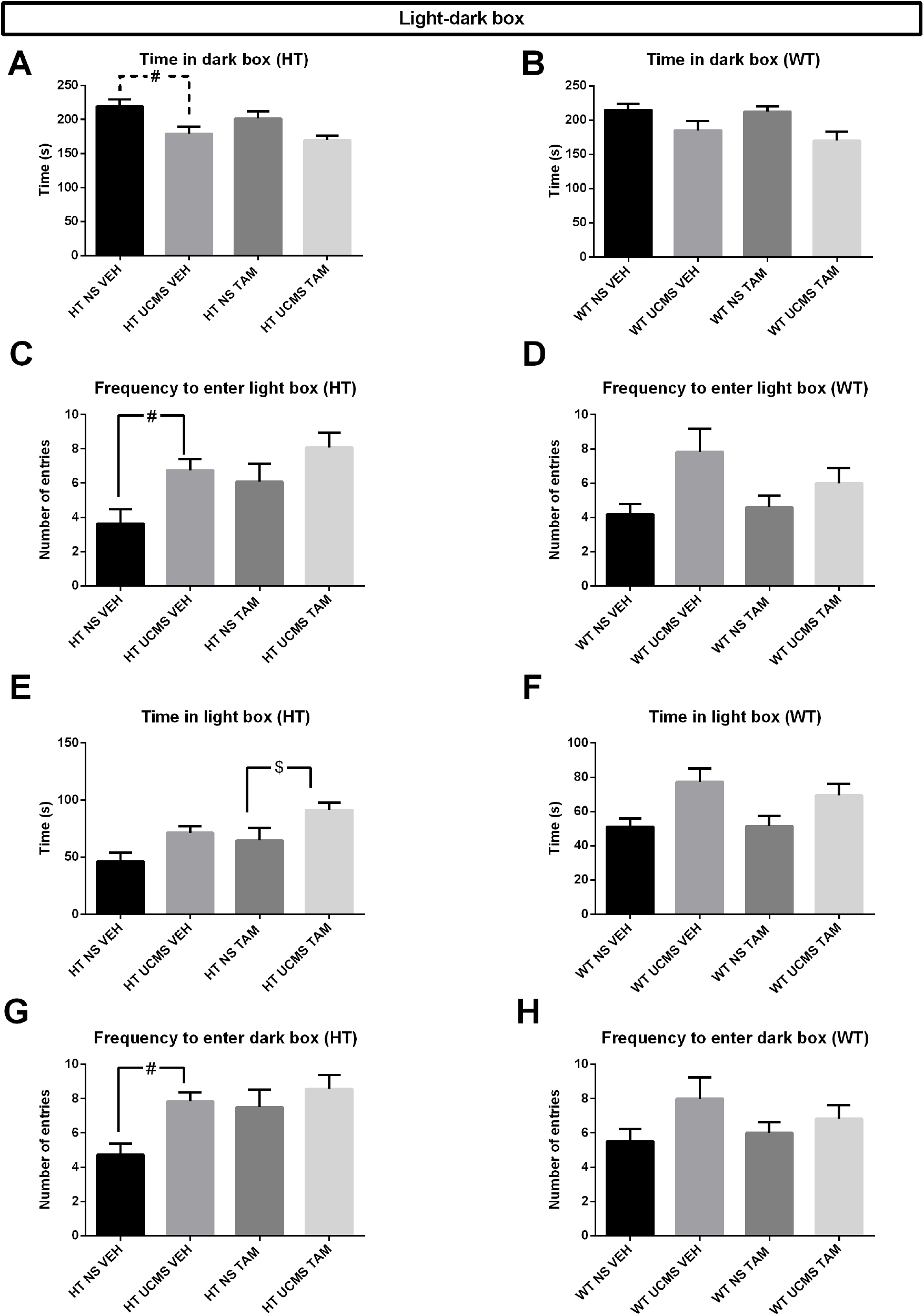
Exposure to UCMS in VEH-treated HT mice increased their frequency of entering both the dark and the light box, while exposure to UCMS in TAM-treated HT mice increased the time spent in the light box. **(A)** In HT animals treated with VEH, exposure to UCMS induced a tendency of decreasing time spent in the dark box. Exposure to UCMS in VEH-treated animals increased the frequency of entering both the **(C)** light box and **(G)** the dark box. **(E)** Exposure to UCMS increased the time spent in the light box for TAM-treated HT animals, n = 11-12 mice per group. **(B, D, F, H)** We did not observe any (a priori chosen) significant pairwise differences between groups of WT animals, n = 10-12 mice per group. Data represent mean ±SEM. $ P < 0.05 for NS TAM vs UCMS TAM; # P < 0.05 for NS VEH vs UCMS VEH; # with dashed lines P = 0.085 for NS VEH vs UCMS VEH

Results from the splash test are shown in Figure 6. Kruskal-Wallis test did not reveal any differences between groups in any of the behavioral measures in the splash test in both HT and WT animals.

**Figure 6.**
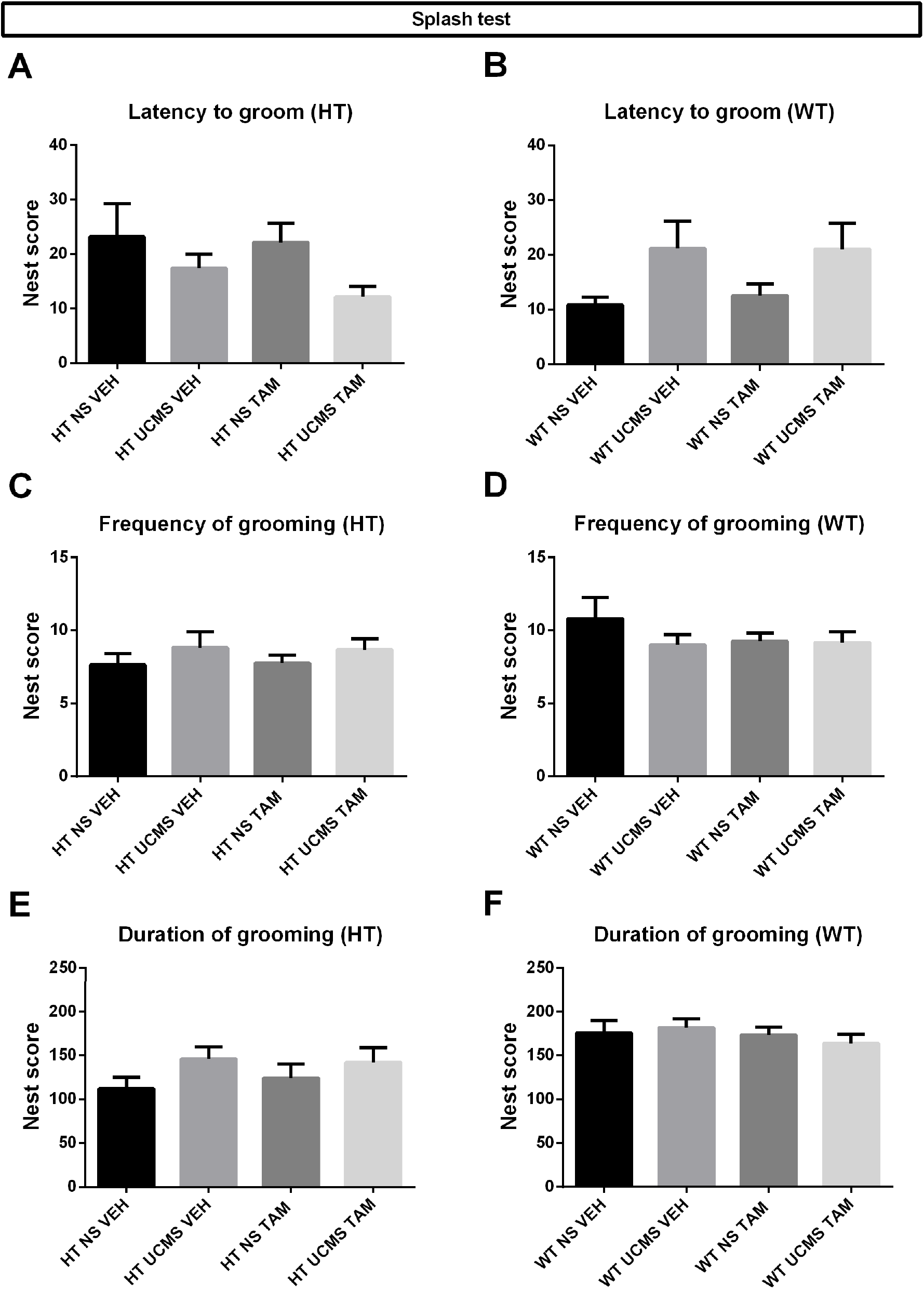
Neither exposure to UCMS nor TAM had any significant effects in the splash test. We did not observe any significant differences between groups in any of the behavioral measures in the splash test in both HT and WT animals. n = 11-12 mice per group. Data represent mean ±SEM.

In the tail suspension test, Kruskal-Wallis test revealed a difference between groups only in HT animals in the time spent immobile between 4 and 6 minutes (*H*_(3,45)_ = 7.89; *P* < 0.05). Post-hoc Dunn’s test revealed only a tendency of tamoxifen to decrease time spent immobile in nonstressed mice (P = 0.078, Figure 7C).

**Figure 7.**
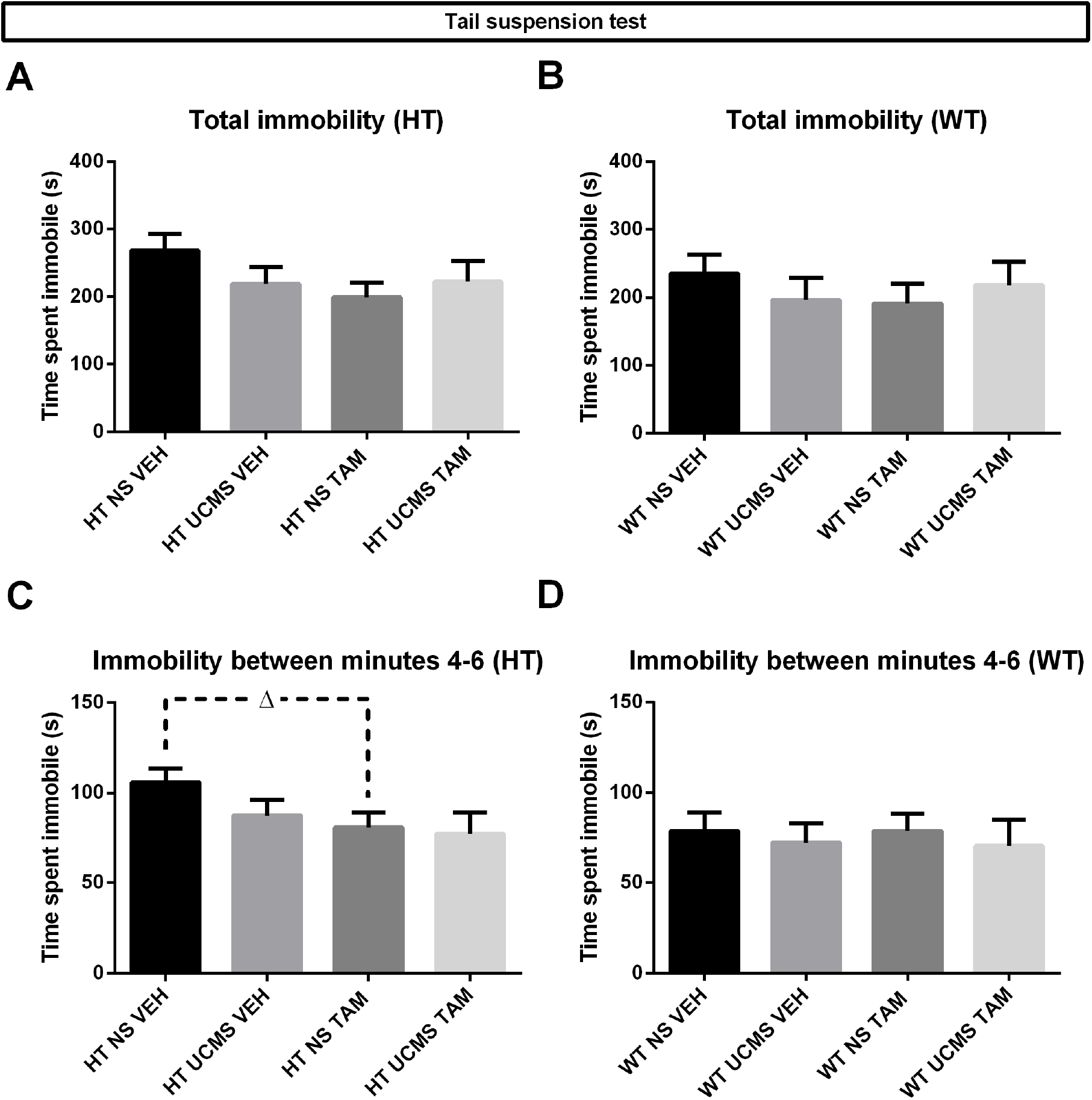
TAM treatment in non-stressed HT mice induced a tendency to decrease immobility time between 4 and 6 minutes in the tail suspension test. **(A)** We did not observe differences between groups of HT mice in total time spent immobile in the tail suspension test, but did observe **(C)** a tendency of tamoxifen to decrease time spent immobile in nonstressed mice between minutes 4 and 6 of the test, n = 11-12 animals per group. **(B, D)** We did not observe any differences between groups in WT animals, n = 10-12 animals per group. Data represent mean ±SEM. Δ with dashed lines P = 0.078 for NS VEH vs NS TAM

In the cookie test, Kruskal-Wallis test revealed a difference between groups only in HT animals in the number of times the animal ate the food in the third session of the cookie test (H_(3,45)_ = 11.57; P < 0.01) and in number of times the animal smelled the food after eating it in the same session (H_(3,45)_ = 11.92; P < 0.01). However, Dunn’s post-hoc test revealed only nonsignificant pairwise differences between groups (i.e., HT NS VEH vs HT UCMS TAM, Figure 8, not designated on the graph).

**Figure 8.**
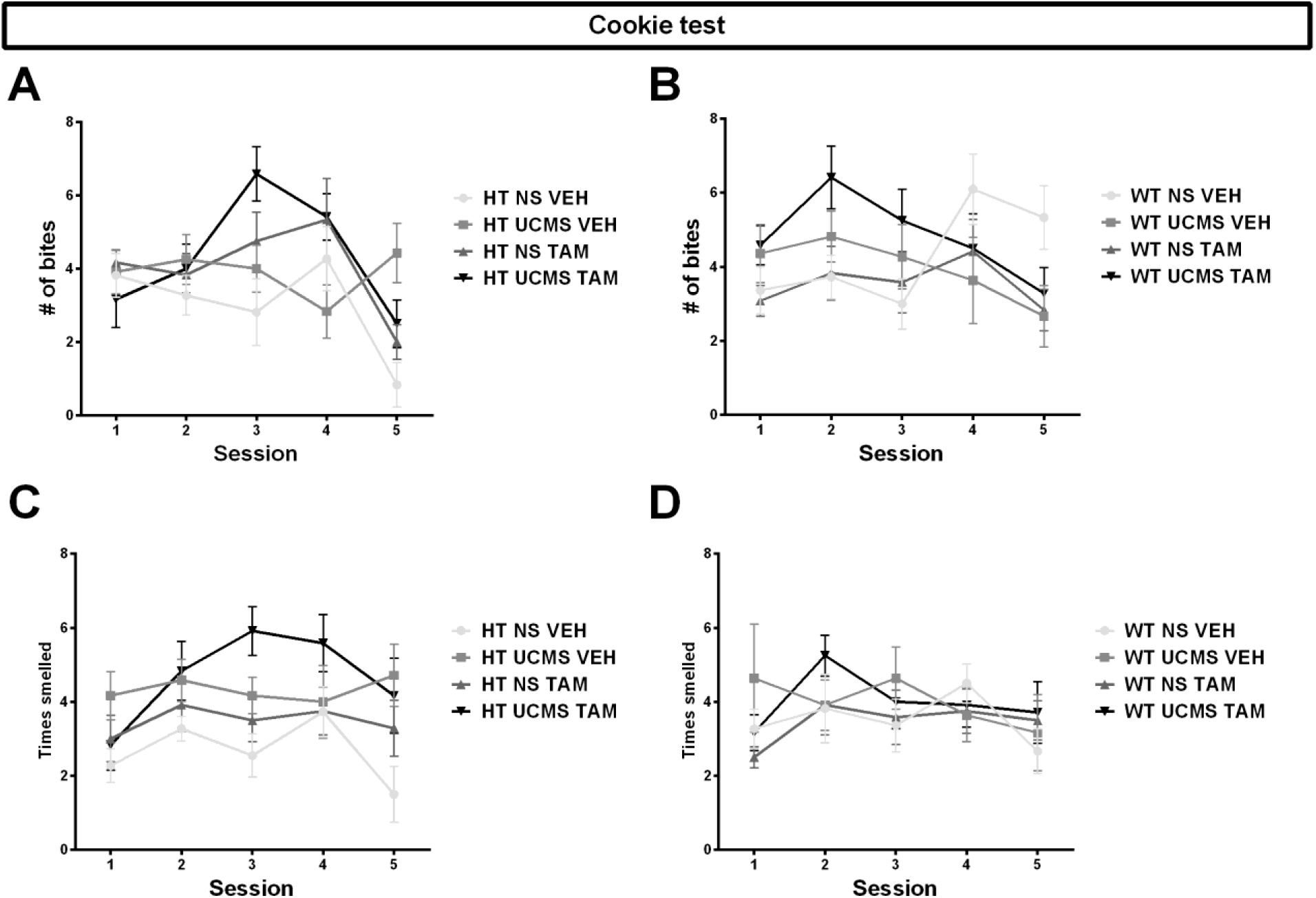
Exposure to UCMS and TAM did not have any significant effects on the number of bites or the times the animal smelled the food in the cookie test. **(A-D)** We did not observe any significant effects of either UCMS or TAM treatment on the number of times the animal ate the food or the number of times the animal smelled the food after first eating it n = 10–12 mice per group. Data represent mean ±SEM.

In the novelty-suppressed feeding test, Kruskal-Wallis test revealed no differences between groups in WT animals, and differences between groups in HT animals in the frequency of smelling the food (H_(3,46)_ = 11.08; P < 0.05), latency to eat (H_(3,46)_ = 8.33; P < 0.05) and in homecage consumption (H_(3,46)_ = 14.29; P < 0.01). Post-hoc Dunn’s tests revealed no (a priori chosen) significant pairwise differences in the frequency of smelling, but did reveal that TAM treatment in mice exposed to stress increased the latency to eat (P < 0.05, Figure 9E) and that exposure to stress increased home-cage consumption in TAM-treated mice (P < 0.05, Figure 9G).

**Figure 9.**
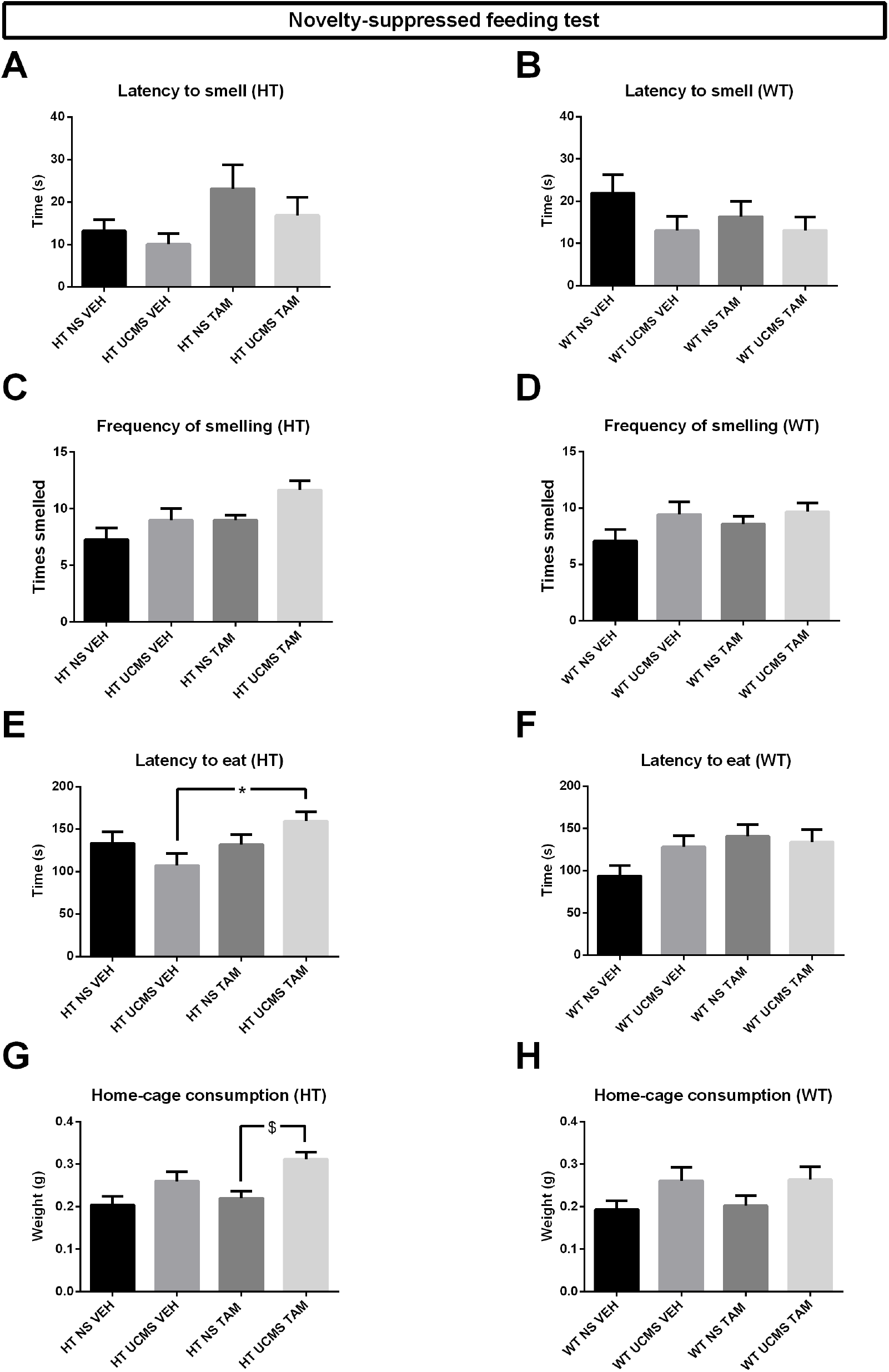
Treatment with TAM increased the latency to eat of HT mice exposed to stress and exposure to stress increased the home-cage consumption of HT mice treated with TAM. **(E)** In HT animals, TAM treatment in mice exposed to stress increased the latency to eat and **(G)** exposure to stress increased home-cage consumption in TAM-treated mice, n = 11-12 animals per group. **(B, D, F, H)** We did not find any significant differences between groups in WT animals, n = 10–12 animals per group. Data represent mean ±SEM. $ P < 0.05 for NS TAM vs UCMS TAM; * P < 0.05 for UCMS VEH vs UCMS TAM

### 3.3. Stress induces a decrease in basal corticosterone levels in TAM-treated mice

To investigate the contribution of adult-born hippocampal neurons on the stress response within the HPA axis, we collected blood from the animals in order to obtain their basal plasma corticosterone levels. We detected a significant difference between groups in basal corticosterone levels in both HT (*H*_(3,46)_ = 12.33; *P* < 0.01) and WT animals (*H*_(3,45)_ = 13.26; P < 0.01). Exposure to stress was able to decrease basal corticosterone levels in TAM-treated mice in both HT (P < 0.01, Figure 10A) and WT animals (P < 0.05, Figure 10B).

**Figure 10.**
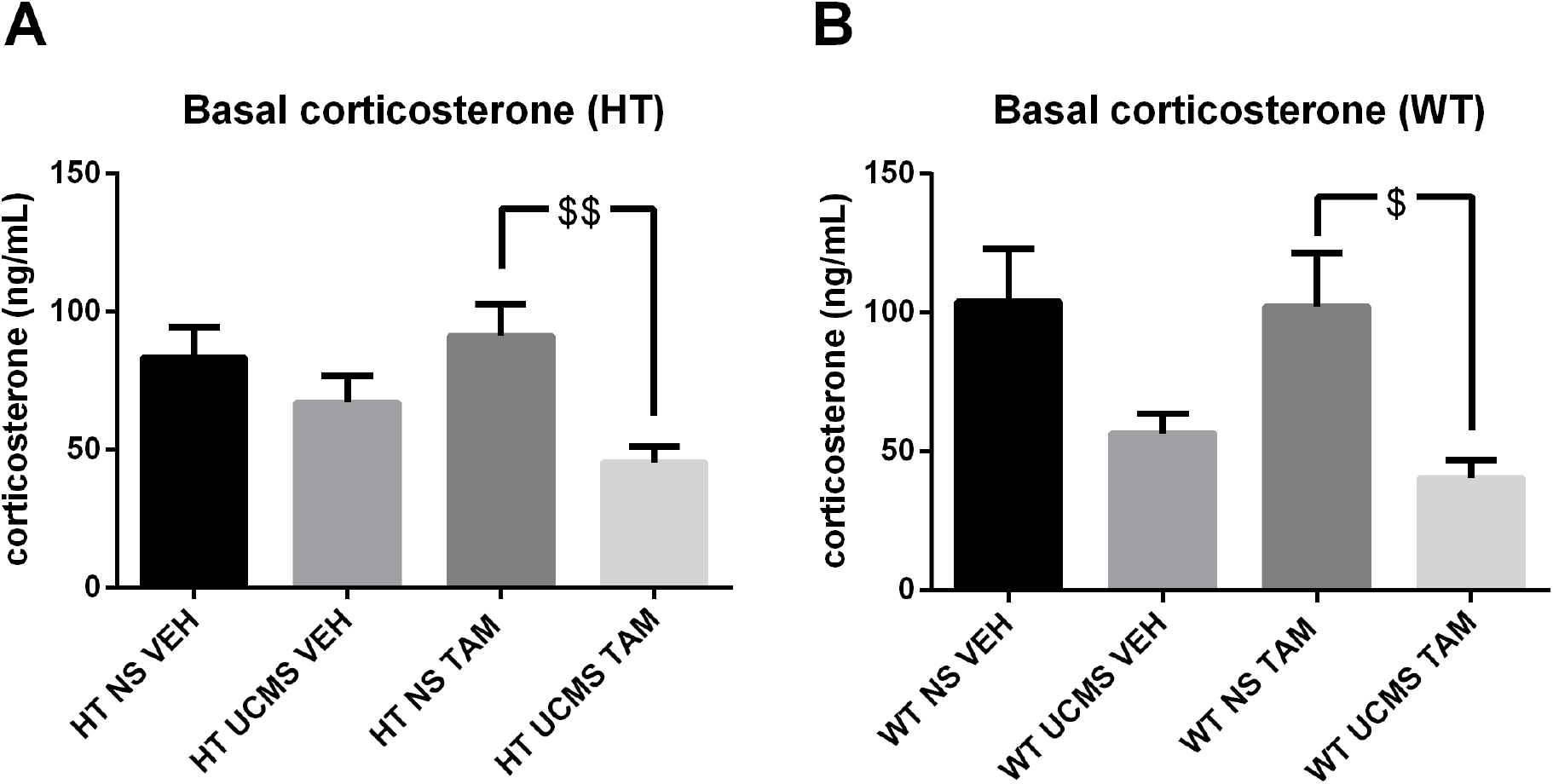
Exposure to UCMS decreased basal corticosterone levels in both HT and WT TAM-treated mice. Exposure to stress was able to decrease basal corticosterone levels in TAM-treated mice in both **(A)** HT and **(B)** WT animals Data represent mean ± SEM. $ P < 0.05 for NS TAM vs UCMS TAM; $$ P < 0.01 for NS TAM vs UCMS TAM

### 3.4. Functional connectivity analysis

We computed a complete set of inter-regional correlations between nodes present in all groups of HT mice, and 66 out of 80 regions of interest met this criterion (Supplementary Table 1 and Figure 11). Network graphs for each group were subsequently generated by considering only the strongest correlations by using a cost threshold with an r-value cut-off calculated to keep the top 10% of nodes in each network. We focused only on the positive correlations between nodes when generating the networks. In all resulting undirected graphs, nodes (brain regions) are connected by edges, which represent inter-regional correlations above the chosen threshold. The correlation matrices varied across groups, with the UCMS VEH group displaying negative correlations between nodes 31-33 (lateral entorhinal cortex, lateral hypothalamic area and lateral posterior thalamic nucleus) with the hippocampal structures, while such negative correlations were not found in other groups (Figure 11A). Graph structure analysis revealed a strong thalamic community in the UCMS groups, which is not as strong in the NS groups (Figure 11B). The NS VEH mice network shows strong connections between the hippocampus and neocortex, while the NS TAM mice network reveals a very strong neocortical community that doesn’t interact much with anything else. The UCMS VEH mice network has strong connections throughout, with the hippocampal community being well connected to the cerebral nuclei community. Both the core and the shell of the NAc are a part of same community as the DG, and they all have high degree centrality. In the UCMS TAM mice network, the hippocampus forms very weak connections, and the strongest connections are between nodes in the cerebral nuclei module and the thalamic module. However, the DG and the paraventricular thalamic nucleus are a part of same community.

**Figure 11.**
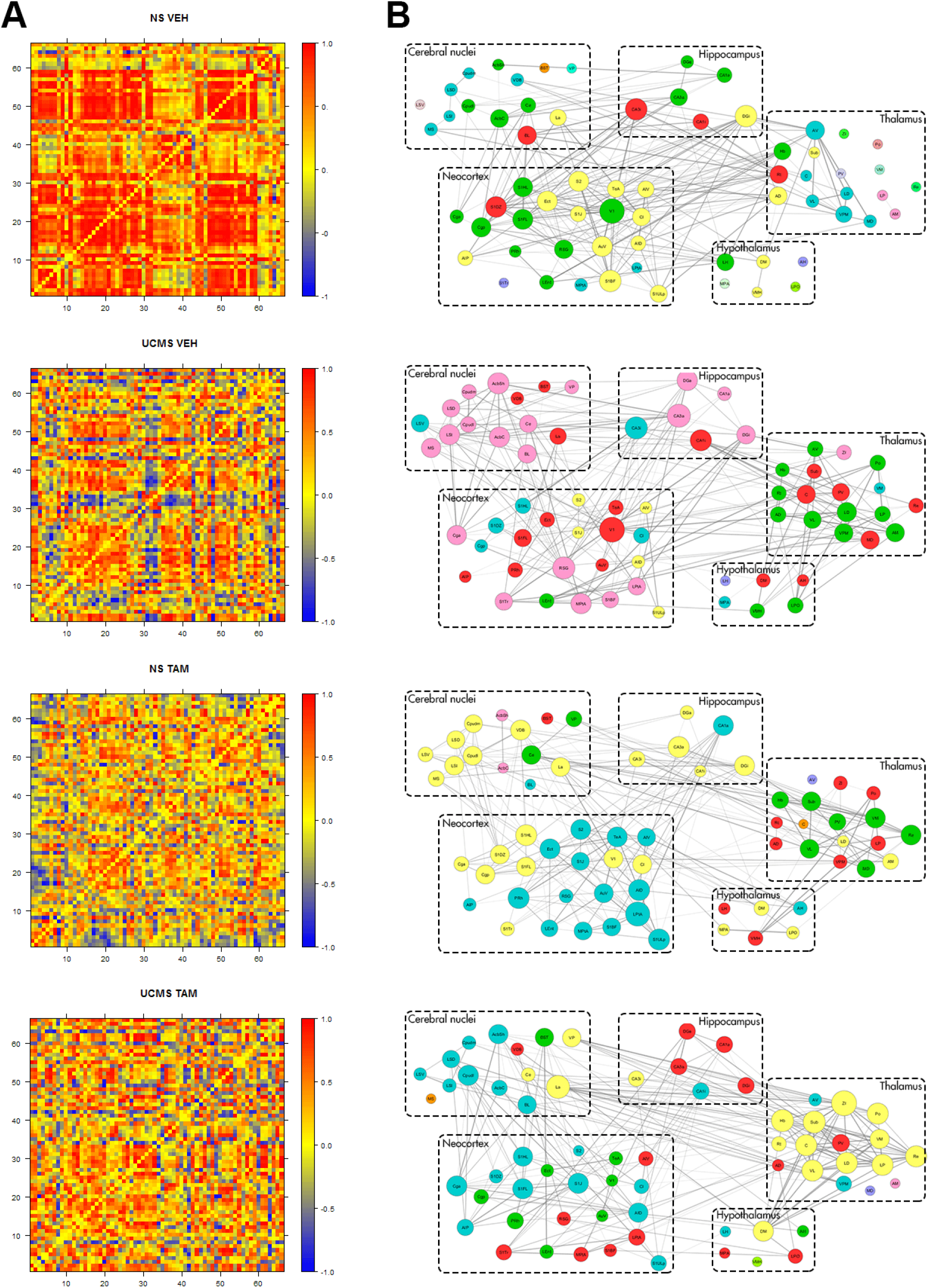
Generation of networks in HT mice. **(A)** Matrices showing inter-regional correlations for ΔFosB expression in 4 groups of HT animals: NS VEH, UCMS VEH, NS TAM and UCMS TAM. Axes are numbered, and correspond to brain regions listed in Supplementary Table 2. Colors reflect correlation strength. **(B)** Network graphs were generated by considering only the strongest correlations (through cost thresholding). In these graphs, regions are grouped by major brain subdivision and node size is proportional to its degree (i.e., number of links per node), while the weight of the connection is proportional to correlation strength. The color of the node corresponds to the community the node belongs to (i.e., nodes with the same color belong to the same community). Abbreviations for brain areas are listed in Supplementary Table 2.

To evaluate whether the generated networks have a small-world organization, we generated control networks with random topology (matched for node, degree and degree distribution) for each network. We quantified the small-worldness by a small-world coefficient, σ, calculated by dividing γ (transitivity of the graph divided by the transitivity of the random graph) with λ (global efficiency of the random graph divided by the global efficiency of the graph), which can be denoted by:

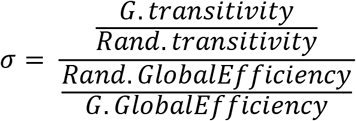

The values of σ for graphs of all groups were calculated to be the following: *σ_NSVEH_* = 1.89; *σ_UCMSVEH_* = 2.77; *σ_NSTAM_* = 2.64 and *σ_UCMSTAM_* = 2.85. Since all of them have a value of σ over 1, that indicates that all the generated networks exhibit small-world topology. Global efficiency is the average inverse shortest path length in the network (Latora and Marchiori, 2001). Transitivity is a measure of segregation, which allows for specialized processing in more densely-connected clusters (Wheeler et al., 2013). It is a weighted version of the clustering coefficient that is less biased by low degree nodes (Rubinov and Sporns, 2010).

For each network, we then identified highly-connected regions (or hubs), which may disproportionately influence network function (Figures 12–15). First we ranked all nodes by degree, which describes the number of links per node, and considered nodes highly-connected if they ranked above the 90^th^ percentile in the network. We then categorized each node to one of five groups by major brain subdivision: cerebral nuclei, hippocampus, neocortex, thalamus and hypothalamus (Figure 16). Because of how the node interacts with the rest of the network depends also on the nature of these connections, we ranked nodes by another measure of centrality – betweenness – which computes the number of shortest paths between node pairs that pass through a given node, and nodes that have a large number of intermodular connections tend to have high betweenness centrality (Sporns et al., 2007; Wheeler et al., 2013). Similarly to the ranking by degree, we considered nodes highly-connected if they ranked above the 90th percentile in the network, and categorized them by major brain subdivision (Figure 17). The nodes which satisfied both criteria (>90^th^ percentile rank) were considered hubs (Figure 18).

**Figure 12.**
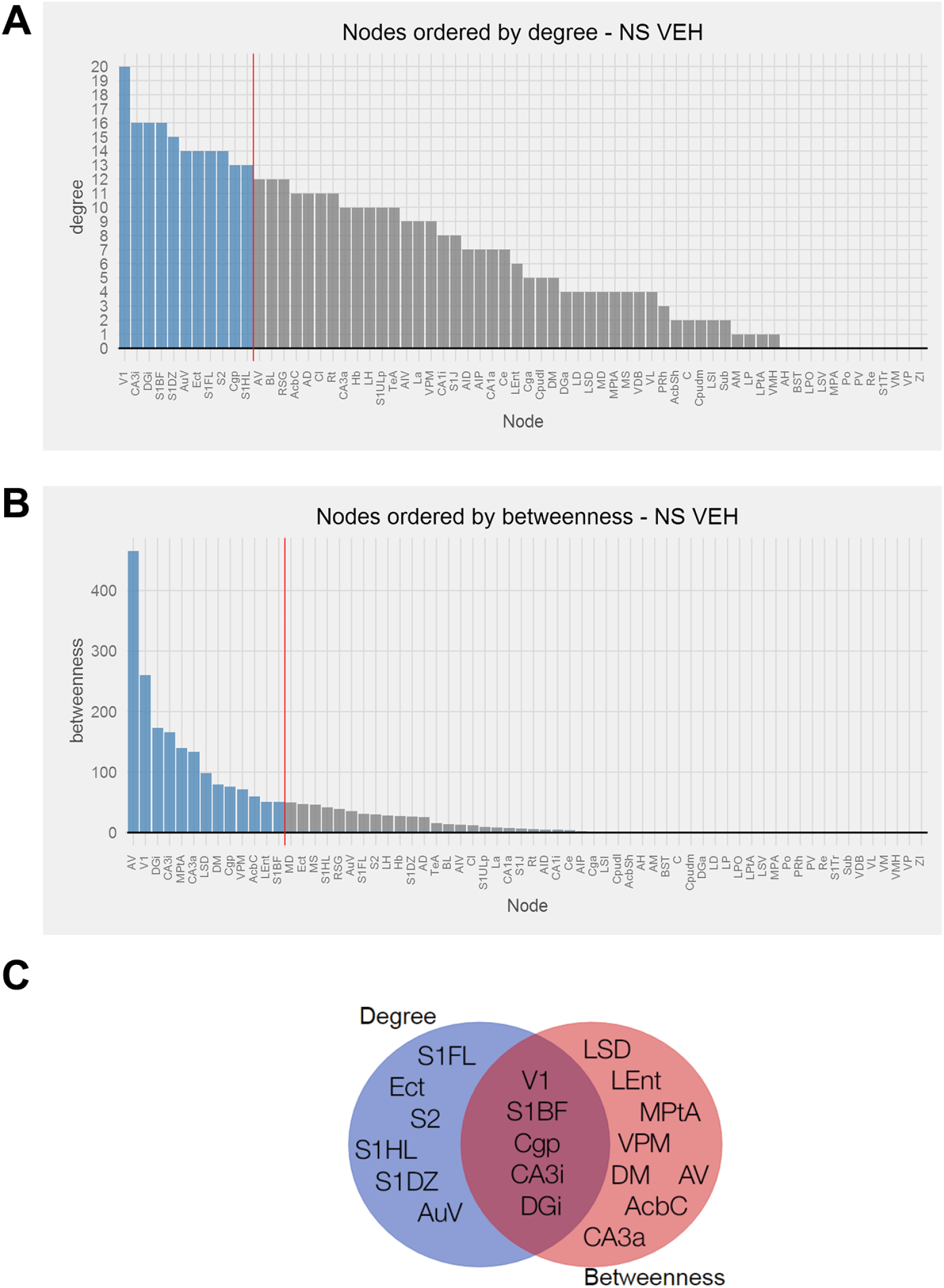
Identification of hubs in the NS VEH generated network. **(A)** Brain regions ranked in descending order by degree and **(B)** betweenness. Vertical red line designates the cut-off point between regions above and below the 90^th^ percentile. **(C)** The overlap between brain regions ranked above the 90^th^ percentile for degree and betweenness in the form of a Venn diagram. Abbreviations for brain areas are listed in Supplementary Table 2.

**Figure 13.**
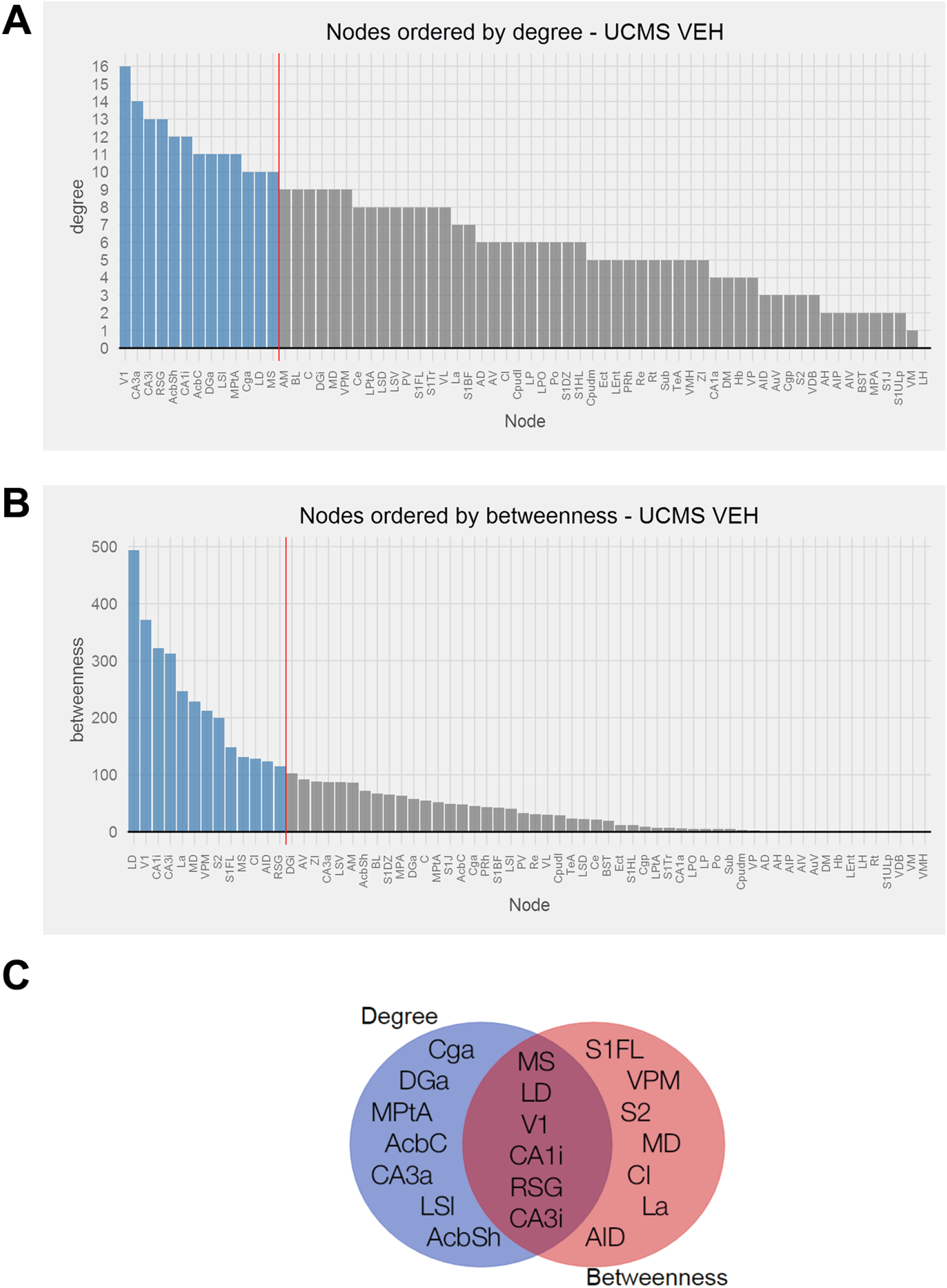
Identification of hubs in the UCMS VEH generated network. **(A)** Brain regions ranked in descending order by degree and **(B)** betweenness. Vertical red line designates the cut-off point between regions above and below the 90^th^ percentile. **(C)** The overlap between brain regions ranked above the 90^th^ percentile for degree and betweenness in the form of a Venn diagram. Abbreviations for brain areas are listed in Supplementary Table 2.

**Figure 14.**
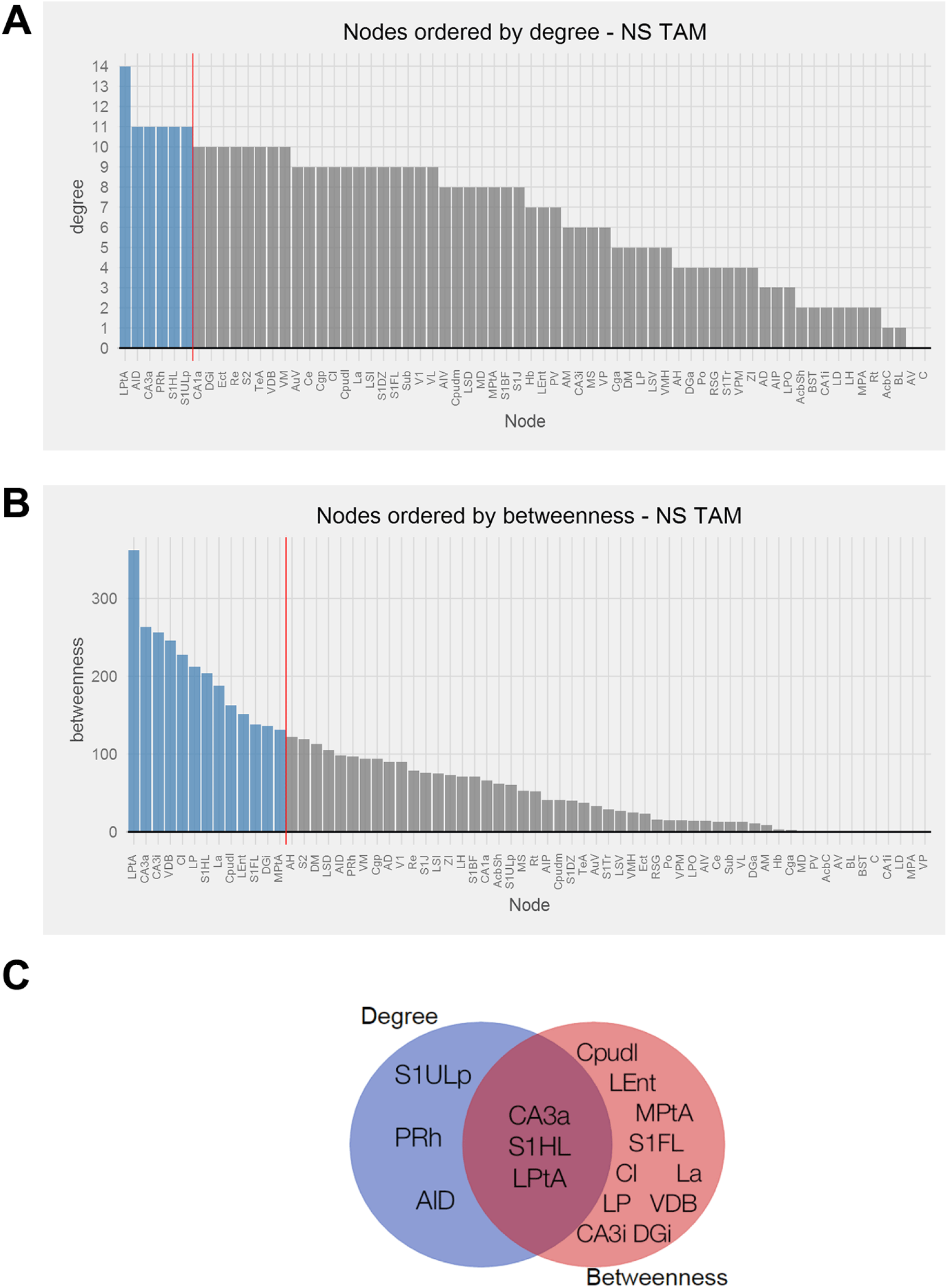
Identification of hubs in the NS TAM generated network. **(A)** Brain regions ranked in descending order by degree and **(B)** betweenness. Vertical red line designates the cut-off point between regions above and below the 90^th^ percentile. **(C)** The overlap between brain regions ranked above the 90^th^ percentile for degree and betweenness in the form of a Venn diagram. Abbreviations for brain areas are listed in Supplementary Table 2.

**Figure 15.**
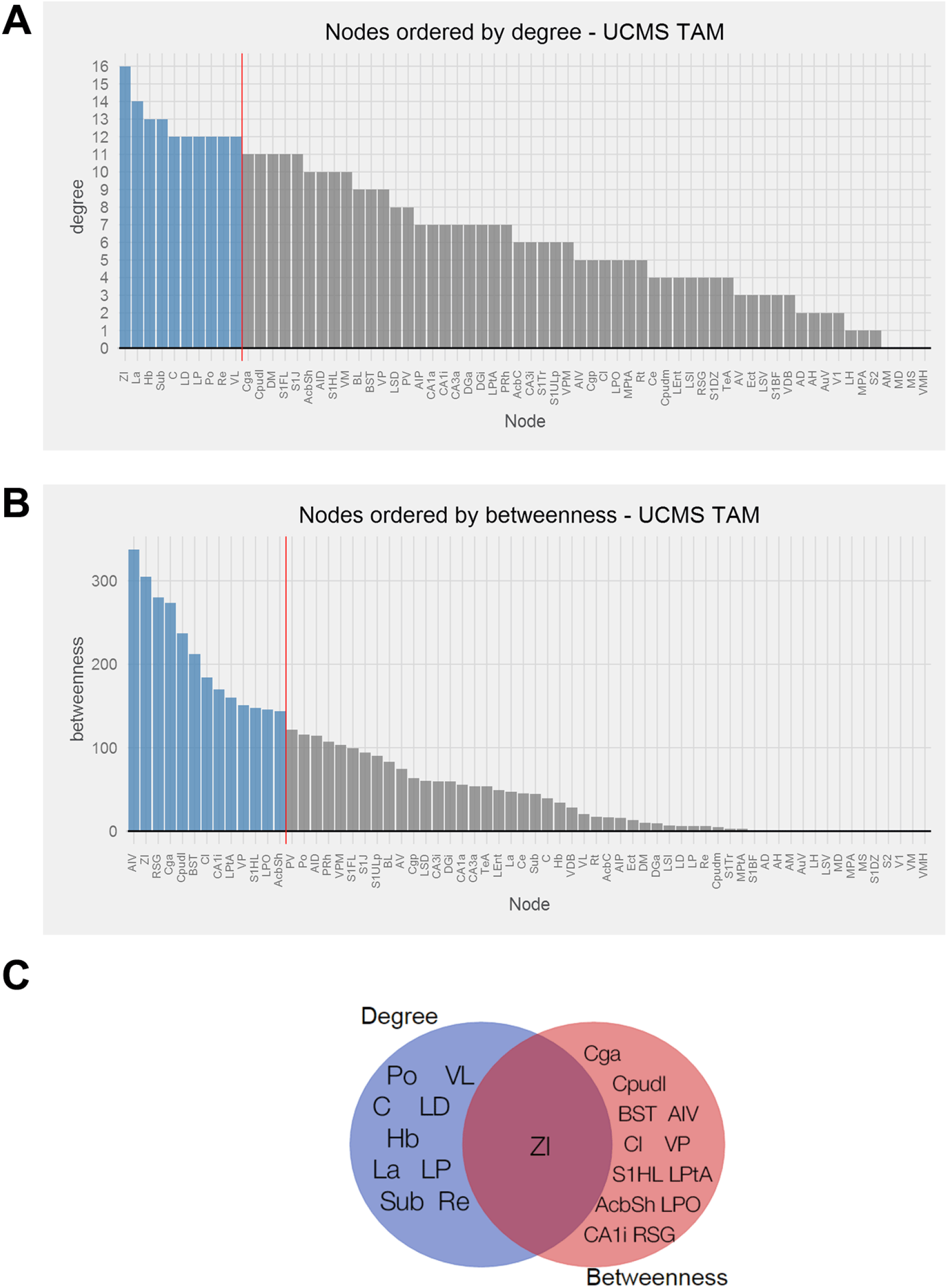
Identification of hubs in the UCMS TAM generated network. **(A)** Brain regions ranked in descending order by degree and **(B)** betweenness. Vertical red line designates the cut-off point between regions above and below the 90^th^ percentile. **(C)** The overlap between brain regions ranked above the 90^th^ percentile for degree and betweenness in the form of a Venn diagram. Abbreviations for brain areas are listed in Supplementary Table 2.

**Figure 16.**
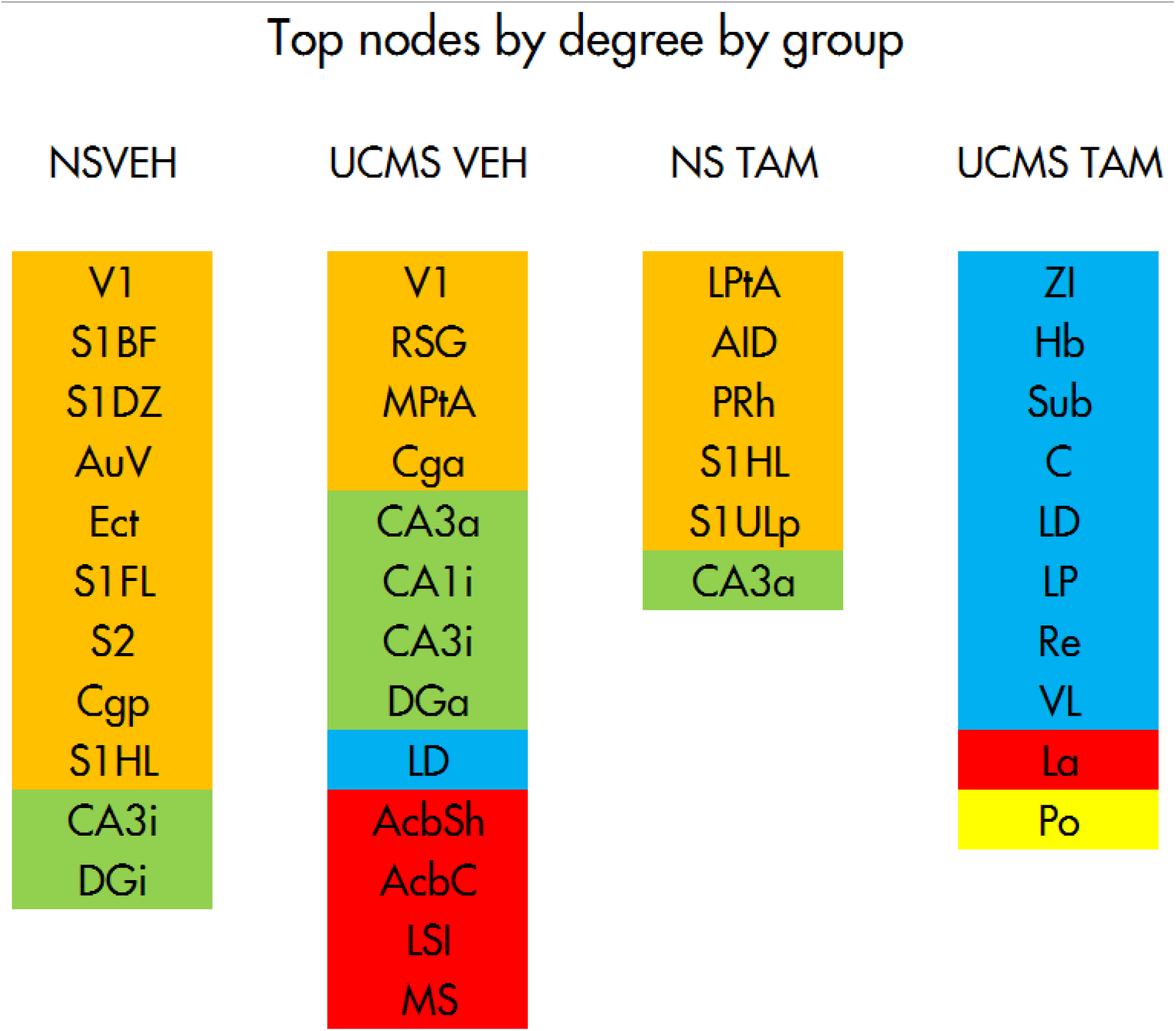
Summary of nodes above the 90th percentile for degree in each group. Brain regions ranked above the 90^th^ percentile for degree with the color of the region corresponding to the major brain division it belongs to. Orange: neocortex; green: hippocampus; blue: thalamus; yellow: hypothalamus and red: cerebral nuclei. Abbreviations for brain areas are listed in Supplementary Table 2.

**Figure 17.**
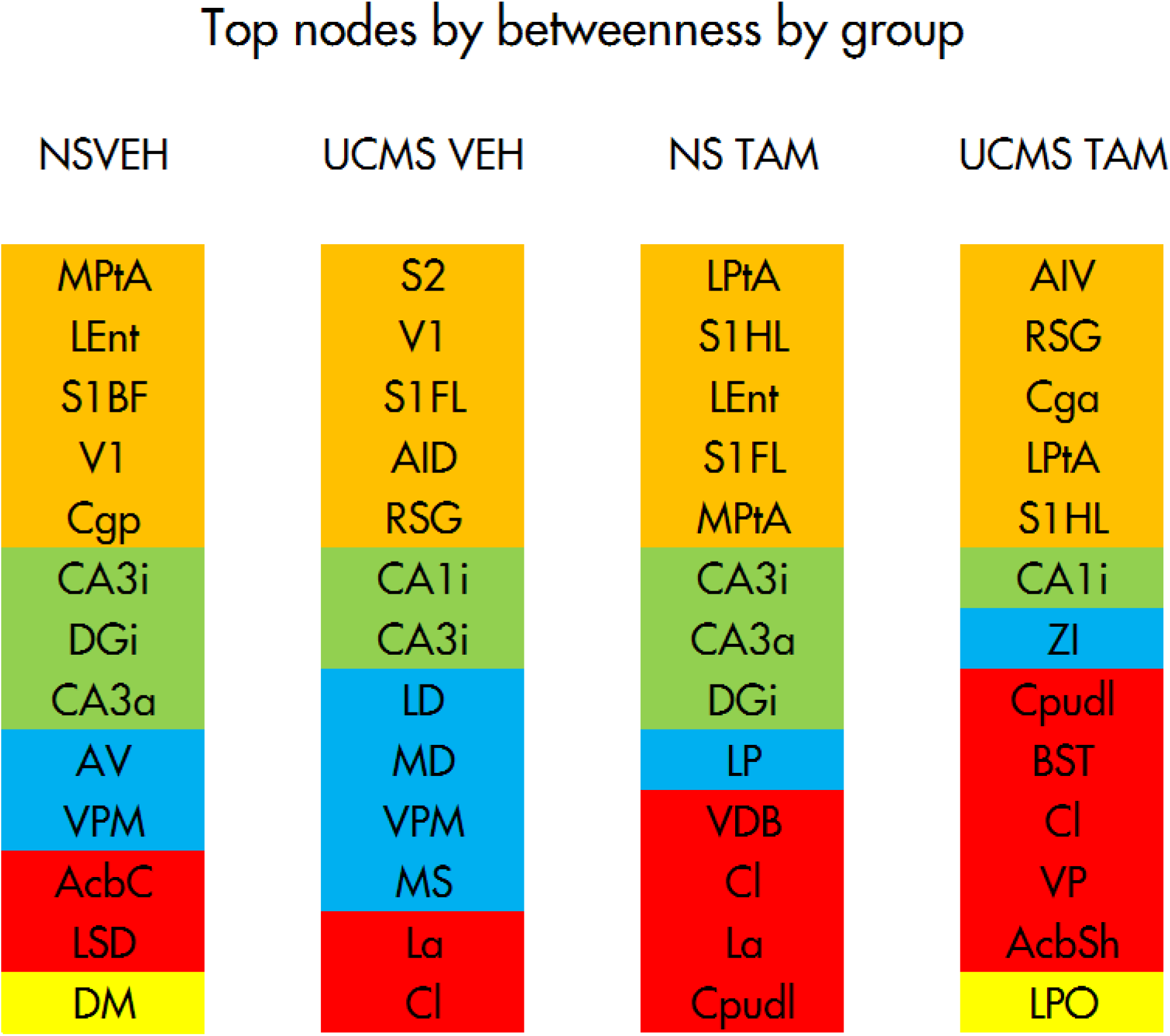
Summary of nodes above the 90th percentile for betweenness in each group. Brain regions ranked above the 90^th^ percentile for betweenness with the color of the region corresponding to the major brain division it belongs to. Orange: neocortex; green: hippocampus; blue: thalamus; yellow: hypothalamus and red: cerebral nuclei. Abbreviations for brain areas are listed in Supplementary Table 2.

**Figure 18.**
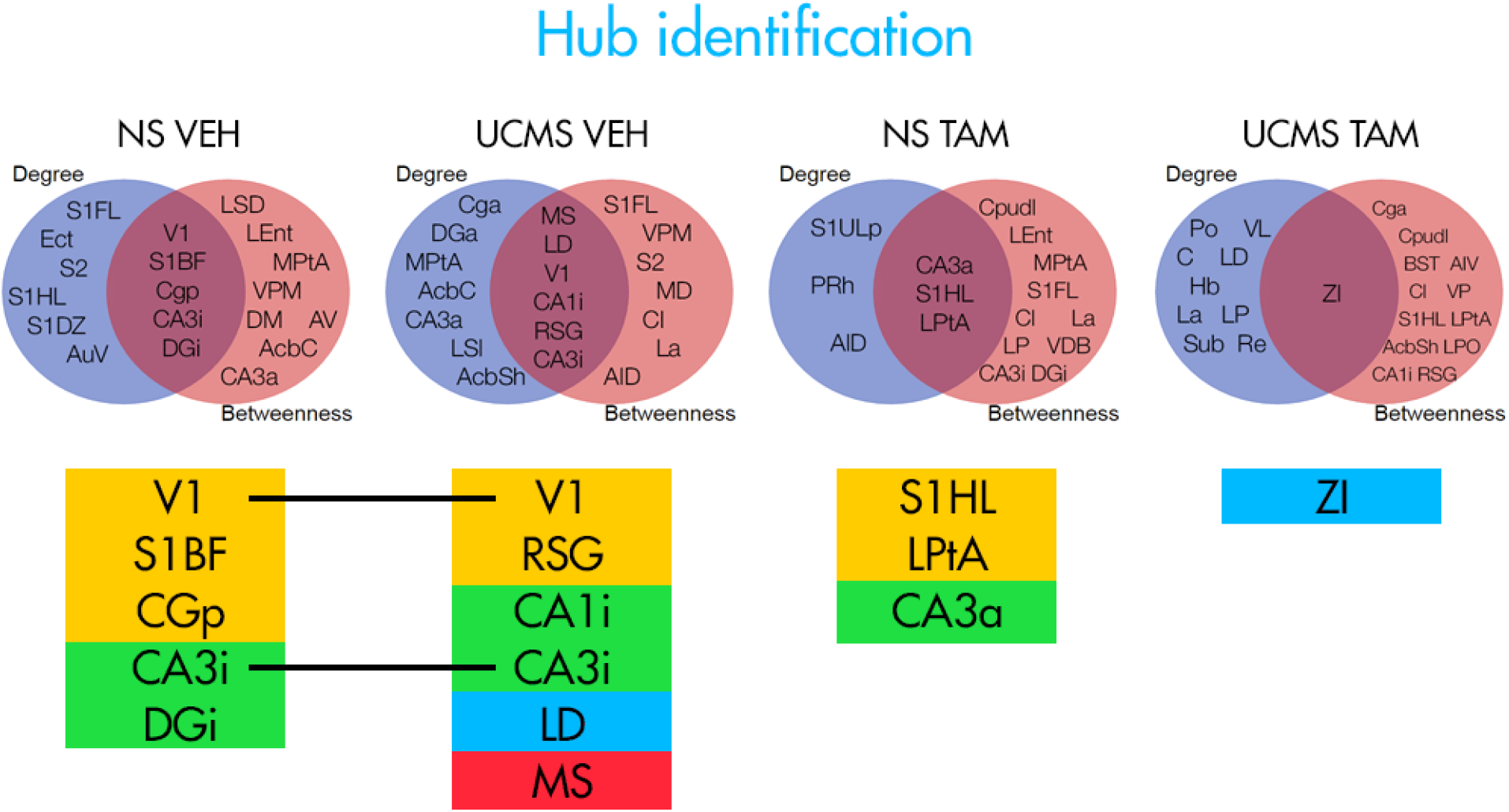
Summary of hub identification across all groups. Brain regions ranked above the 90^th^ percentile for both degree and betweenness were considered hubs. The color of the region corresponds to the major brain division it belongs to. Orange: neocortex; green: hippocampus; blue: thalamus and red: cerebral nuclei. Abbreviations for brain areas are listed in Supplementary Table 2.

### 3.5. Western blot

To assess the specificity of the antibody used for immunohistochemistry, western blots were performed on micro-dissected tissue from all groups of animals. The analysis of western blots is in progress.

## 4. Discussion

There have been recent advancements in linking the manipulations of adult neurogenesis to antidepressant-like effects, which were achieved by utilizing a transgenic mouse model that enables researchers to explore the effects of specifically increasing neurogenesis while not affecting other biological processes. This approach has several advantages over other (non- or less-specific) approaches such as environmental enrichment, voluntary exercise and administration of the P7C3 compound, which in addition to increasing neurogenesis also affect other biological processes (Hendriksen et al., 2012; Latchney et al., 2015; Llorens-Martin et al., 2011; Wang et al., 2014). The main advantage is the ability to dissociate the specific effects of an increase in neurogenesis from all the other effects, enabling researchers to identify a strict causal link between neurogenesis and behavior. To the best of our knowledge, only two studies so far have explored this link in an animal model of depression, with one study increasing neurogenesis before chronic corticosterone administration (Hill et al., 2015), and the other one increasing neurogenesis after the onset of unpredictable chronic mild stress (Culig et al., 2017). The first study showed that increasing neurogenesis before exposure to an insult is able to decrease vulnerability against same insult, while in the second study we showed that increasing neurogenesis after the onset of stress can elicit partial remission. However, it is still an open question through which mechanism is the small number of adult-born neurons in the dentate gyrus able to mediate a behavioral and endocrine response to stress.

We attempted to extend these findings by extensive behavioral phenotyping and functional network construction through graph theory analysis, tools enabling us to identify changes in longterm functional connectivity after exposure to stress (i.e., “the stressome”) and how a specific increase in neurogenesis might mediate remission. We exposed the animals to an insult (UCMS) and then attempted to increase neurogenesis during the same challenging conditions, which is a paradigm that mimics the timeline of depression onset and treatment in humans. Two weeks of exposure to UCMS in TAM-treated HT mice was enough to cause a significant deterioration in the coat state, showing that a depressive-like phenotype was induced, which is consistent with our observations from the previous experiment (Culig et al., 2017).

We assessed the levels of neurogenesis in the DG by immunostaining for the immature neuronal marker DCX, like in previous experiments utilizing *iBax* mice (Culig et al., 2017; Hill et al., 2015; Sahay et al., 2011). Unlike in the previous experiments, we show that administration of TAM did not increase the number of DCX+ cells in the DG and that UCMS did not decrease their number. Therefore, the inability of TAM to increase the levels of neurogenesis beyond the basal levels (and of UCMS to decrease them) suggests that the behavioral and physiological differences observed cannot be attributed to a change in neurogenesis levels. One potential explanation of this lack of effect lies in the age of the animals; when we divided the animals in two distinct subsets (“younger” (13-14 weeks at behavioral testing) and “older” (18-22 weeks at behavioral testing) animals) and analyzed neurogenesis in separate age groups, we found no effects present in the younger cohort of mice (Supplementary Figure 1A), but found that in the older cohort of mice (Supplementary Figure 1B), TAM treatment induced a significant increase of the density of DCX+ cells in the DG of UCMS animals, which is a result that partially reproduces what we observed in our previous study (Culig et al., 2017). Therefore, one possible explanation is that the basal levels of neurogenesis in the younger cohort of mice were too high for either UCMS to decrease them or TAM to increase them (ceiling effect), and that by mixing these two distinct age groups, we were unable to observe any effects on neurogenesis.

On the physical level, we observed the adverse effect of UCMS on the coat state, consistent with previous findings (Ibarguen-Vargas et al., 2008). We failed to replicate the previous finding of tamoxifen improving the coat state of animals in Week 9 (Culig et al., 2017), and we attribute this to the failure of tamoxifen to induce an increase in neurogenesis in all animals. We do, however, observe a trend of TAM decreasing the coat state scores in HT UCMS mice (compared to its matched VEH group), but the lack of statistical significance might be a result of a reduced sample size (10-12 animals per group, as opposed to 17-26 per group in the previous experiment). It would also be worthwhile to examine the coat state score dynamics in a longer UCMS protocol, as TAM seems to produce observable effects after a 6 week long lag after administration, correlating well with the fact that newborn neurons might contribute to behavior when aged 4-6 weeks, possibly through enhanced synaptic plasticity with increased LTP amplitude and decreased LTP induction threshold (Song et al., 2012).

Exposure to UCMS induced effects in several behavioral readouts of HT mice. We replicated our previous findings that exposure to stress is able to increase the scores in the nest building test at the +5h time point in TAM-treated mice (Culig et al., 2017). While this effect of stress seems paradoxical and warrants further investigation, we propose that this effect might be a consequence of NS mice having to adapt to a novel environment (an acute stressor), while UCMS mice were transferred in a bigger cage with cotton, corresponding to an improvement to their condition, enabling them to construct the nest faster. However, as this effect was not observed in VEH-treated mice, or in WT animals, the result seems to be a specific consequence of TAM administration, which is surprising given the fact that we failed to observe an effect of TAM on neurogenesis. Similarly, exposure to UCMS also induced an increase in the time spent in the light box in the light-dark box, which is usually interpreted as either an increase in exploratory behavior or as an anxiolytic effect (Bourin and Hascoёt, 2003). A similar paradoxical effect has been observed in our previous experiment, where exposure to UCMS decreased the amount of time spent in the dark box (Culig et al., 2017), and we interpreted this result similarly to the nest building test results, as a specific effect of TAM in HT mice exposed to stress. Additionally, exposure to UCMS increased the home-cage consumption of HT TAM-treated mice in the NSF test, making the other behavioral readouts from this test difficult to appreciate, as differences in behavior could be a result of differences in appetite or food-related reward (Stedenfeld et al., 2011). Interestingly, in the NSF test, we found that TAM treatment in mice exposed to UCMS increased the latency to eat the food (despite HT UCMS TAM animals having the highest homecage consumption), which is a readout that is usually interpreted as an increase in anxiety-like behavior (Dulawa and Hen, 2005; Surget et al., 2008). Another surprising effect of exposure to UCMS was a decrease in basal corticosterone levels for TAM-treated mice. However, this is consistent with the results from our previous experiment (Culig et al., 2017), and a possible explanation might be that the mice exposed to UCMS were already accustomed to frequent handling and various stressors (while the NS mice were not) making it possible that the increased CORT levels in the NS groups are a consequence of intense acute stress of blood sampling. TAM administration seems to have had opposite effect on NS and UCMS mice, with TAM seemingly increasing the basal CORT in HT NS mice, and decreasing it in HT UCMS mice. In further experiments, it would be valuable to use a less invasive method of assessing CORT levels (such as through fecal collection), in order to minimize the effects of exposure to an acute stressor on plasma CORT levels.

In terms of effects of exposure to UCMS in HT VEH mice, we found that exposure to UCMS significantly increased the frequencies to enter the light and the dark box in the light-dark box test, as well as to induce a tendency to decrease time spent in the dark box. Decreased time spent in the dark box is often interpreted as an decrease in anxiety-like behavior (Costall et al., 1989; Young and Johnson, 1991), while increased transitions from the light to the dark box (and vice versa) are usually interpreted as a reduction in anxiety-like behavior (Bourin and Hascoët, 2003; Crawley et al., 1984). Finally, we observed an effect of TAM in HT NS mice in the tail suspension test, where TAM administration decreased the time spent immobile between the 4^th^ and the 6^th^ minute of the test in the NS condition, a readout which is usually interpreted as a decrease of behavioral despair. We did not observe any other significant effects in locomotor activity, minimizing the possible confounding effects of a difference in baseline locomotor activity on other behavioral tests. Also, no effects were observed in the cookie test, which is a result that complements our findings from the previous experiment, where an increase in neurogenesis was not able to modulate hedonic behavior in the sucrose preference test (Culig et al., 2017). Not observing even an effect of UCMS exposure on hedonic behavior might be a consequence of decreased sensitivity to stress of *iBax* mice.

In regards to the ΔFosB expression and inter-regional correlations, we observed UCMS-induced negative correlations between the lateral entorhinal cortex and both the hippocampal structures and the nucleus accumbens (shell and core) in the VEH-treated mice, which were not present in other groups. When ranking nodes by their degree, we observe a strong thalamic-cortical signature in both NS groups. Exposure to UCMS seems to induce activity in thalamic areas and cerebral nuclei, with a different signature in the UCMS TAM group, which seems to completely “disengage” the neocortex and has most of its top nodes in the thalamus. The UCMS VEH group engages different areas (the shell and the core of the NAc) in the “cerebral nuclei” group of nodes, and UCMS TAM engages the lateral amygdaloid nucleus (La). However, one shortcoming of this analysis is a different number of nodes identified in the top 90^th^ percentile for degree between groups (e.g., 6 in NS TAM and 13 in UCMS VEH). This is likely a consequence of the more discrete nature of the degree distribution and should be corrected in further analyses, so that each group contains the same number of nodes above the chosen percentile. We found similar results in hub identification, which identified mostly different brain areas in all groups, but also a disengagement of the neocortical nodes and engagement of the thalamic nodes as a response to exposure to UCMS. Interestingly, the only hub identified in the UCMS TAM group is the Zona Incerta (ZI), a brain area with roles in central pain, compulsions and Parkinson’s disease (Mallet et al., 2002; Masri et al., 2009) that has only recently been shown to modulate fear generalization (Venkataraman et al., 2019). In the graph analysis, we found that our constructed networks of all groups have a small-world topology, like many real-world networks. However, the specific nature of functional connections between nodes depends on both the treatment and the environment the animal group has been exposed to. The functional network of the UCMS groups seems to have a strong thalamic signature, which is not present in the NS groups, while the NS mice treated with TAM display a very strong neocortical community that does not interact much with other areas. However, it is difficult to appreciate the effect of TAM in both the NS and UCMS conditions, given that the networks constructed contain both age-groups of animals, where we failed to detect an effect of TAM on neurogenesis in the DG. While it might be worthwhile to repeat the connectome analysis after dividing the animals into two distinct subgroups (by age), this poses a challenge in terms of statistical analysis, as the number of animals per each subgroup drops to 3, making it difficult to generate correlations and establish sufficient statistical power.

Overall, we were not able to detect an increase in hippocampal neurogenesis as a result of our TAM treatment, making it difficult to analyze the observed subset of results that seem to be a result of TAM treatment. Additionally, exposure to UCMS seems to have affected the mice in only a few behavioral readouts, which could be a consequence of the strain’s higher resilience to our stress protocol, prompting for a less mild UCMS protocol in future studies. Despite these shortcomings, we were able to replicate some of the findings from our previous experiment (Culig et al., 2017), which include paradoxical effects of stress improving the nest building scores and lowering basal corticosterone levels (in TAM-treated mice). Our network analyses identified various hubs across groups, areas which might be of particular importance in influencing overall network function, with Zona Incerta being of particular interest in the UCMS TAM group. Additionally, we were able to detect a strong thalamic signature as a result of stress exposure, which is interesting in the light of certain studies detecting an increased activation of the thalamus in depression (Fu et al., 2004; Kumari et al., 2003). Analyzing both the behavior and the connectome of age-separated groups (where we observe an effect of tamoxifen on neurogenesis) should provide more insight in regards to the connectomic mechanism of remission and, in particular, the role of the DG in modulating the previously observed partial remission from stress-induced depressive-like behavior.

