## Supplementary Materials for "Changes in the functional connectome and behavior after exposure to chronic stress and increasing neurogenesis"

### Supplementary material

| Region | Abbreviation | # |
|---|---|---|
| **Neocortex** | | |
| cingulate cortex - anteror | Cg-a | 19 |
| cingulate cortex - posterior | Cg-p | 20 |
| retrosplenial granular cortex | RSG | 47 |
| ectorhinal cortex | Ect | 27 |
| lateral entorhinal cortex | LEnt | 31 |
| lateral parietal association cortex | LPtA | 35 |
| medial parietal association cortex | MPtA | 41 |
| perirhinal cortex | PRh | 44 |
| temporal association cortex | TeA | 58 |
| secondary auditory cortex, ventral area | AuV | 9 |
| primary visual cortex | V1 | 59 |
| agranular insular cortex, posterior part | AIP | 6 |
| agranular insular cortex, dorsal part | AID | 5 |
| agranular insular cortex, ventral part | AIV | 7 |
| primary somatosensory cortex, barrel field | S1BF | 49 |
| primary somatosensory cortex, dysgranular region | S1DZ | 50 |
| primary somatosensory cortex, forelimb region | S1FL | 51 |
| primary somatosensory cortex, hindlimb region | S1HL | 52 |
| primary somatosensory cortex, jaw region | S1J | 53 |
| primary somatosensory cortex, trunk region | S1Tr | 54 |
| primary somatosensory cortex, upper lip region | S1ULp | 55 |
| secondary somatosensory cortex | S2 | 56 |
| **Thalamus** | | |
| anterodorsal thalamic nucleus | AD | 3 |
| anteromedial thalamic nucleus | AM | 8 |
| anteroventral thalamic nucleus | AV | 10 |
| central thalamic nucleus | C | 13 |
| habenular nucleus | Hb | 28 |
| laterodorsal thalamic nucleus | LD | 30 |
| lateral posterior thalamic nucleus | LP | 33 |

| | | |
|---|---|---|
| mediodorsal thalamic nucleus | MD | 39 |
| posterior thalamic nuclear group | Po | 43 |
| reuniens thalamic nucleus | Re | 46 |
| submedius thalamic nucleus | Sub | 57 |
| ventrolateral thalamic nucleus | VL | 61 |
| ventromedial thalamic nucleus | VM | 62 |
| ventral posteromedial thalamic nucleus | VPM | 65 |
| reticular thalamic nucleus | Rt | 48 |
| zona inserta | ZI | 66 |

**Hypothalamus**

| | | |
|---|---|---|
| anterior hypothalamic area | AH | 4 |
| dorsomedial hypothalamic nucleus | DM | 26 |
| ventromedial hypothalamic nucleus | VMH | 63 |
| lateral hypothalamic area | LH | 32 |
| lateral preoptic area | LPO | 34 |
| medial preoptic area | MPA | 40 |

**Cerebral nuclei**

| | | |
|---|---|---|
| bed nucleus of the stria terminalis | BST | 12 |
| caudate putamen (striatum) - dorsal lateral | Cpu-dl | 22 |
| caudate putamen (striatum) - dorsal medial | Cpu-dm | 23 |
| accumbens nucleus, core | AcbC | 1 |
| accumbens nucleus, shell | AcbSh | 2 |
| ventral pallidum | VP | 64 |
| nucleus of the vertical limb of the, diagonal band | VDB | 60 |
| claustrum | Cl | 21 |
| lateral amygdaloid nucleus | La | 29 |
| central amygdaloid nucleus | Ce | 18 |
| basolateral amygdaloid nucleus | BL | 11 |
| medial septal nucleus | MS | 42 |
| lateral septal nucleus, dorsal part | LSD | 36 |
| lateral septal nucleus, intermediate part | LSI | 37 |
| lateral septal nucleus, ventral part | LSV | 38 |

| | | |
|---|---|---|
| **Hippocampus** | | |
| field CA1 of hippocampus - anterior | CA1-a | 14 |
| field CA1 of hippocampus - intermediate | CA1-i | 15 |
| field CA3 of hippocampus - anterior | CA3-a | 16 |
| field CA3 of hippocampus - intermediate | CA3-i | 17 |
| dentate gyrus - anterior | DG-a | 24 |
| dentate gyrus - intermediate | DG-i | 25 |

**Supplementary Table 1. List of brain regions in which ΔFosB was quantified with corresponding abbreviations and reference numbers for the correlation matrices. Brain regions are categorized by major brain subdivision.**

| Img | label | manual_loose | auto_count | perror | difference |
|---|---|---|---|---|---|
| 298B2_07 | 19 | 682 | 758 | 11,14% | 5,28% |
| 298B2_07 | 20 | 710 | 864 | 21,69% | 9,78% |
| 298B2_07 | 21 | 555 | 874 | 57,48% | 22,32% |
| 298B2_07 | 22 | 367 | 581 | 58,31% | 22,57% |
| 298B2_07 | 25 | 1179 | 1738 | 47,41% | 19,16% |
| 298B2_07 | 26 | 1256 | 1443 | 14,89% | 6,93% |
| 298B2_07 | 27 | 525 | 752 | 43,24% | 17,78% |
| 298B2_07 | 28 | 390 | 582 | 49,23% | 19,75% |
| 298B2_07 | 29 | 402 | 598 | 48,76% | 19,60% |
| 298B2_07 | 30 | 289 | 406 | 40,48% | 16,83% |
| 298B2_07 | 31 | 576 | 759 | 31,77% | 13,71% |
| 298B2_07 | 32 | 684 | 938 | 37,13% | 15,66% |
| 298B2_07 | 33 | 1929 | 2261 | 17,21% | 7,92% |
| 298B2_07 | 34 | 1571 | 2143 | 36,41% | 15,40% |
| 298B2_07 | 67 | 87 | 43 | 50,57% | 33,85% |
| 298B2_07 | 68 | 77 | 28 | 63,64% | 46,67% |
| 298B2_07 | 69 | **13** | 4 | 69,23% | 52,94% |
| 298B2_07 | 70 | 28 | 19 | 32,14% | 19,15% |
| 298B2_07 | 73 | **10** | 4 | 60,00% | 42,86% |
| 298B2_07 | 74 | **7** | 0 | 100,00% | 100,00% |
| 298B2_07 | 77 | **8** | 2 | 75,00% | 60,00% |
| 298B2_07 | 78 | **6** | 2 | 66,67% | 50,00% |
| 298B2_07 | 79 | 68 | 43 | 36,76% | 22,52% |
| 298B2_07 | 80 | 75 | 54 | 28,00% | 16,28% |
| 298B2_07 | 89 | **172** | 6 | 96,51% | 93,26% |
| 298B2_07 | 90 | **177** | 10 | 94,35% | 89,30% |
| 298B2_07 | 91 | 45 | 19 | 57,78% | 40,63% |
| 298B2_07 | 92 | 37 | 28 | 24,32% | 13,85% |
| 298B2_07 | 99 | **66** | 3 | 95,45% | 91,30% |
| 298B2_07 | 100 | **114** | 10 | 91,23% | 83,87% |
| 298B2_07 | 105 | 25 | 17 | 32,00% | 19,05% |
| 298B2_07 | 106 | 34 | 37 | 8,82% | 4,23% |
| 298B2_07 | 137 | 114 | 159 | 39,47% | 16,48% |
| 298B2_07 | 138 | 118 | 161 | 36,44% | 15,41% |
| 298B2_07 | 147 | 446 | 827 | 85,43% | 29,93% |
| 298B2_07 | 148 | 461 | 876 | 90,02% | 31,04% |
| 298B2_07 | 153 | 261 | 349 | 33,72% | 14,43% |
| 298B2_07 | 154 | 193 | 299 | 54,92% | 21,54% |
| 298B2_07 | 159 | 639 | 1026 | 60,56% | 23,24% |
| 298B2_07 | 160 | 753 | 1065 | 41,43% | 17,16% |

| | | | | | |
|---|---|---|---|---|---|
| 270B1_06 | 39 | 482 | 516 | 7,05% | 3,41% |
| 270B1_06 | 40 | 296 | 307 | 3,72% | 1,82% |
| 270B1_06 | 41 | 390 | 429 | 10,00% | 4,76% |
| 270B1_06 | 42 | 239 | 127 | 46,86% | 30,60% |
| 270B1_06 | 51 | 801 | 598 | 25,34% | 14,51% |
| 270B1_06 | 52 | 578 | 486 | 15,92% | 8,65% |
| 270B1_06 | 57 | 793 | 661 | 16,65% | 9,08% |
| 270B1_06 | 58 | 601 | 531 | 11,65% | 6,18% |
| 270B1_06 | 123 | 1323 | 1716 | 29,71% | 12,93% |
| 270B1_06 | 124 | 1484 | 1758 | 18,46% | 8,45% |
| 270B1_06 | 125 | 1370 | 1588 | 15,91% | 7,37% |
| 270B1_06 | 126 | 1165 | 1545 | 32,62% | 14,02% |
| 270B1_06 | 129 | 144 | 215 | 49,31% | 19,78% |
| 270B1_06 | 130 | 147 | 229 | 55,78% | 21,81% |
| | | | **average** | 24,21% | 11,67% |
| | | | **stdev** | 25,99% | 24,62% |

**Supplementary Table 2. Comparison of automated cell identification with manual counts from an experimenter**

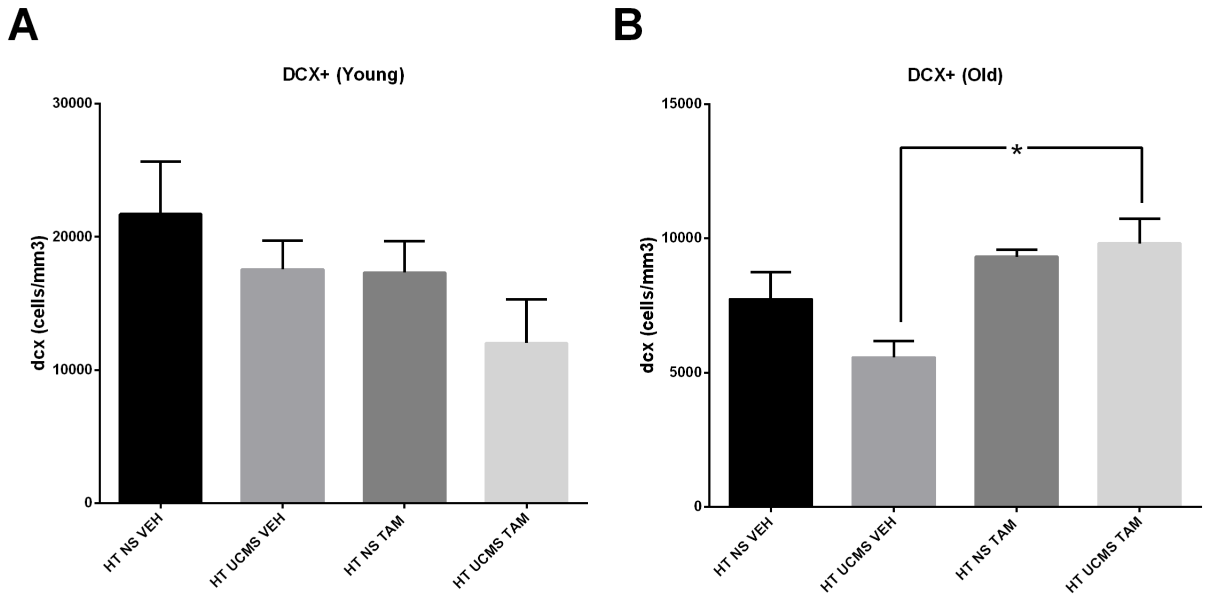


**Supplementary Figure 1. Treatment with TAM only had an effect on neurogenesis in the older cohort of HT animals subjected to UCMS**

**(A)** Neither exposure to UCMS nor TAM administration had an effect on the number of DCX+ cells in the DG of the younger cohort of mice. **(B)** Two-way ANOVA revealed a significant effect of treatment in the older cohort of mice, and post-hoc Bonferroni's multiple comparison test detected that TAM treatment induced a significant increase of the density of DCX+ cells in the DG of UCMS animals, n = 2–4 per group. Data represent mean ± SEM. * $P < 0.05$ UCMS VEH vs UCMS TAM

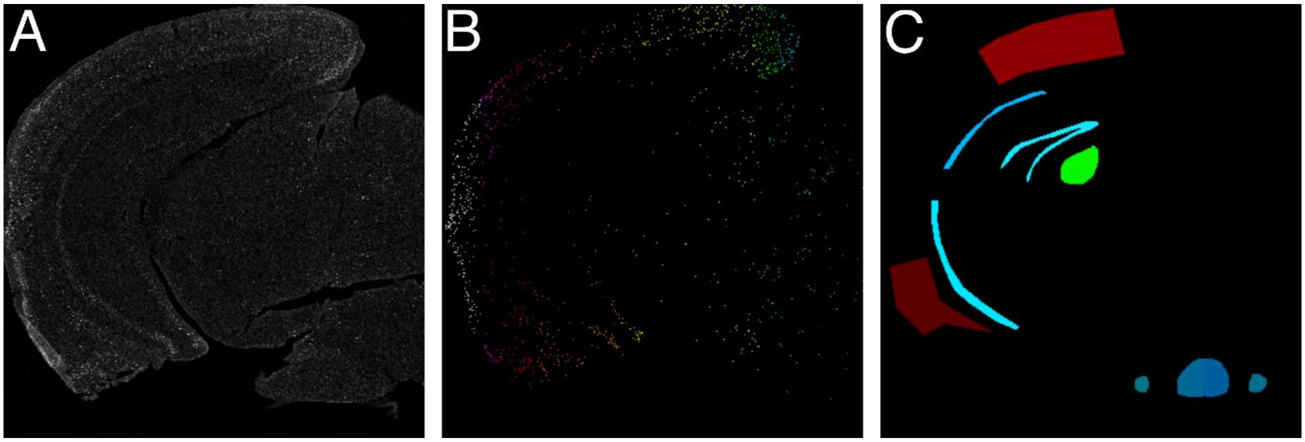


**Supplementary Figure 2. Automated cell counting in brain ROIs**

**(A)** Brain section after initial image manipulations. **(B)** Visualization of cells detected by the algorithm (rainbow colors used just for visualization purposes). **(C)** A mask where image density counts will be calculated, every color represents a different ROI.
